# STING-Activating Polymer-Drug Conjugates for Cancer Immunotherapy

**DOI:** 10.1101/2024.03.23.585817

**Authors:** Taylor L. Sheehy, Alexander J. Kwiatkowski, Karan Arora, Blaise R. Kimmel, Jacob A. Schulman, Katherine N. Gibson-Corley, John T. Wilson

## Abstract

The stimulator of interferon genes (STING) pathway links innate and adaptive antitumor immunity and therefore plays an important role in cancer immune surveillance. This has prompted widespread development of STING agonists for cancer immunotherapy, but pharmacological barriers continue to limit the clinical impact of STING agonists and motivate the development of drug delivery systems to improve their efficacy and/or safety. To address these challenges, we developed SAPCon, a STING-activating polymer-drug conjugate platform based on strain-promoted azide-alkyne cycloaddition of dimeric-amidobenzimidazole (diABZI) STING agonists to hydrophilic polymer chains through an enzyme-responsive chemical linker. To synthesize a first-generation SAPCon, we designed a diABZI prodrug modified with a DBCO reactive handle with a cathepsin B-cleavable spacer for intracellular drug release and conjugated this to pendant azide groups on a 100kDa poly(dimethyla acrylamide-*co*-azide methacrylate) copolymer backbone to increase circulation time and enable passive tumor accumulation. We found that intravenously administered SAPCon accumulated at tumor sites, where it was endocytosed by tumor-associated myeloid cells, resulting in increased STING activation in tumor tissue. Consequently, SAPCon promoted an immunogenic tumor microenvironment, characterized by increased frequency of activated macrophages and dendritic cells and improved infiltration of CD8^+^ T cells, resulting in inhibition of tumor growth, prolonged survival, and enhanced response to anti-PD-1 immune checkpoint blockade in orthotopic breast cancer models. Collectively, these studies position SAPCon as a modular and programmable platform for improving the efficacy of systemically administered STING agonists for cancer immunotherapy.

## Introduction

As immunotherapy continues to transform the current cancer treatment landscape, there remains an unmet need for improved and durable outcomes for the ∼15% of patients still minimally responsive to immune checkpoint blockade (ICB) targeting PD-1 or PD-L1.^1^ For many patients, the response to ICB therapy is dependent on the immunological phenotype of the tumor microenvironment (TME).^2^ Although there is clear evidence that ICB can stimulate antitumor immunity, patients with “cold” tumors lack sufficient proinflammatory cell infiltration, such as activated macrophages and dendritic cells, and, most notably, cytotoxic CD8^+^ T cells, which are often required for optimal response.^3, 4, 5^ The presence of these cell types in the TME is important for proper function of the cancer immunity cycle (CIC) and often correlates with improved response to ICB antibodies.^6,7^ Hence, there is a significant demand for therapeutic platforms that enrich the TME with immunological cues that shift it towards a “hot,” proinflammatory and T cell-enriched state.

The cyclic guanosine monophosphate-adenosine monophosphate synthase (cGAS)-stimulator of interferon genes (STING) pathway is a highly conserved, pattern recognition pathway that has been identified as a key mediator of antitumor immunity.^8^ cGAS recognition of cytosolic DNA, a sign of cellular distress and disease, results in the synthesis of the cyclic dinucleotide, 2’3’-cyclic guanosine monophosphate-adenosine monophosphate (2’3’cGAMP) which binds to STING on the cytosolic side of the endoplasmic reticulum.^9^ STING activation promotes expression of type-I interferon (IFN-I) and interferon-stimulated genes (ISGs), which has been attributed to enhanced tumor-antigen specific, cytotoxic T cell responses.^10^ IFN-I promotes processing and presentation of tumor antigens by APCs with consequent activation of CD4^+^ and CD8^+^ T cells. STING activation also results in the production of chemokine gradients (i.e. CXCL9 and CXCL10) that direct T cells, APCs, and natural killer cells to the TME.^11,12^ In fact, functional STING has been shown to be critical for optimal response to ICB in some mouse models.^13^

Due its important role in antitumor immunity, STING pathway agonists are currently being explored as cancer immunotherapies. Cyclic dinucleotide (CDNs) STING agonists exert robust antitumor effects in preclinical mouse models and but have thus far failed to advance in clinical trials.^14^ While multiple factors have likely contributed to these disappointing outcomes, it can be partially attributed to the use of an intratumoral administration route. This is not only impractical for many patients and tumor types but has lacked clinical benefit across a wide spectrum of immunomodulatory agents. However, systemic administration of CDNs remains limited by several drug delivery barriers such as rapid clearance, poor tumor accumulation, and minimal intracellular delivery.^15^ Small molecule STING agonists have been developed to circumvent some of these barriers and offer potential for systemic administration.^16 17 18^ For example, Ramanjulu and co-workers at GlaxoSmithKline developed one such promising small molecule, a dimeric-amidobenzimidazole (diABZI) (Compound 3), with 400-fold more *in vitro* potency compared to 2’3’ cGAMP and a half-life of 1.4 h compared to a few minutes for 2’3’-cGAMP, resulting in antitumor efficacy in CT26 colorectal and B16.F10 melanoma murine models following intravenous administration. ^16^

While diABZI and other STING agonists are currently under clinical investigation (e.g., NCT03843359, NCT04420884, NCT04609579), they lack tumor or cell specificity, and concerns exist regarding the consequences of indiscriminate, systemic STING activation. For example, non-specific, systemic STING activation may result in cytokine release syndrome with potentially adverse consequences that have been described for other immunotherapies such as CAR T cells.^19,20^ Similarly, STING activation has been shown to exacerbate some chronic inflammatory and autoimmune diseases, such as ulcerative colitis, nonalcoholic fatty liver disease, and lupus.^21^ STING agonists can also induce apoptosis in CD8^+^ T cells, which are critical antitumor effectors.^22^ Hence, there is a need to develop drug delivery platforms to improve the pharmacological properties of systemically administered STING agonists and enhance their activity in tumors and/or secondary lymphoid organs.

Polymer-drug conjugates provide a versatile approach to modulating drug pharmacokinetics (PK) and biodistribution (BD) profiles and can be designed to enhance tumor accumulation through passive (e.g., EPR effect) and/or active (e.g., ligand targeted) mechanisms. They can be further engineered to allow for environmentally-responsive drug release at tumor sites or within specific cell populations.^23,24, 25^ While polymer-drug conjugates have been extensively employed for delivery of chemotherapeutics and have advanced to clinical trials^26,27^, there has been only minimal investigation into their design and optimization for delivery of STING agonists.^28,29^ Herein, we present STING-activating polymer-drug conjugates (SAPCon) as a modular platform for systemic administration of STING agonists (**Figure 1**). Our approach leverages an inert, water-soluble poly(dimethyl acrylamide) (DMA) backbone and a “clickable” DBCO-functionalized diABZI STING agonist that can be covalently conjugated to azide-functionalized polymer scaffolds through cathepsin cleavable linkers. We demonstrate that SAPCon offers extended half-life, enhanced tumor accumulation, and enzyme-mediated drug release to increase STING activation at tumor sites, resulting in immunological remodeling of the TME that promotes antitumor immunity and improves response to anti-PD-1 ICB in murine models of breast cancer. This positions SAPCon as a versatile and enabling strategy for improving systemic delivery of STING agonists to enhance cancer immunotherapy.

**Figure 1:**
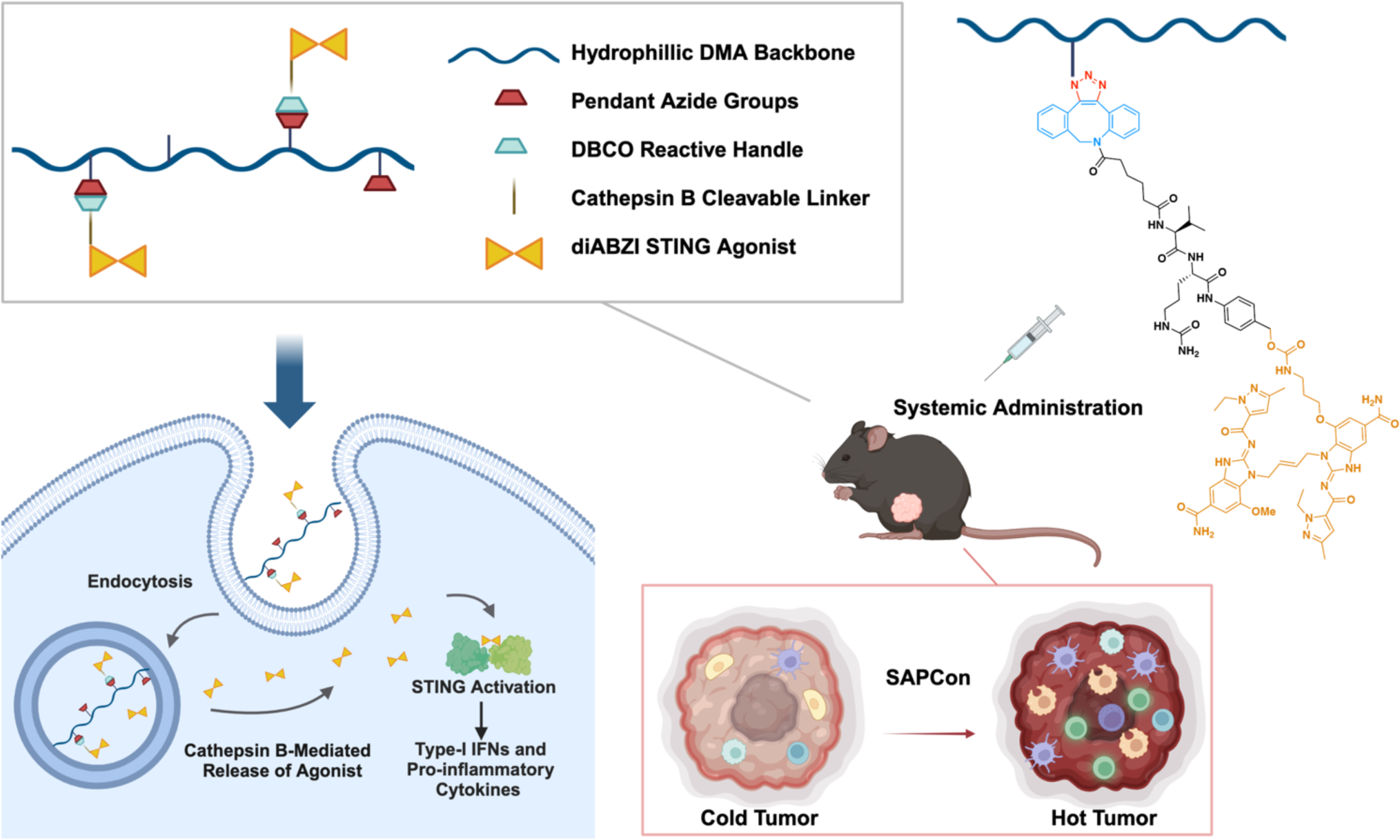
Design of STING-Activating Polymer Drug Conjugates (SAPCon). SAPCon consists of a hydrophilic poly(dimethylacrylamide) (DMA) backbone co-polymerized with azide-functionalized monomers for strain-promoted azide-alkyne cycloaddition reaction to a DBCO-functionalized, cathepsin B-cleavable, diABZI STING agonist. The SAPCon platform can be delivered intravenously to promote tumor accumulation and cellular uptake and enzyme-mediated diABZI release by tumor-associated myeloid cells to enhance STING activation in tumor tissue, resulting in reprogramming of the tumor-immune microenvironment. Figure created using BioRender and ChemDraw 21.

## Results and Discussion

### Design and Characterization of Conjugatable and Enzymatically-Responsive diABZI STING Agonist

Several design principles were considered in conceiving and synthesizing a STING agonist for polymer-mediated delivery. Here, we selected a diABZI-based compound due to its high affinity and immunostimulatory potency, as well as availability of chemically accessible sites outside of the STING binding pocket that allow for modification and introduction of reactive handles. In particular, the 7-position of the benzimidazole is exposed and therefore amenable to modification. We synthesized a diABZI variant with a versatile primary amine group that allows for facile conjugation to diverse drug linkers and other reactive handles. Additionally, the charged amine group increases water solubility and reduces membrane permeability, a strategy that has been deployed for antibody-drug conjugates to reduce activity of drug that is prematurely released from the carrier.^30,31^ The parent diABZI-amine compound **(2)** and relevant intermediates were synthesized as detailed in **Scheme S1**. Building from diABZI-amine, we next installed a reactive handle for ligation to polymer carriers and a cleavable linker to allow for intracellular diABZI release (**Figure 2A**). A self-immolative, cathepsin-cleavable valine-citrulline-PAB linker (denoted V/C henceforth) was chosen due to its common use in polymer and antibody drug conjugates and the potential for tumor-specific drug release due to the upregulation of cathepsins in the TME.^32,33,34^ While a multitude of bioconjugation chemistries could be used to link diABZI to polymeric carriers, we elected for dibenzocyclooctyne (DBCO) handles that can undergo strain-promoted azide-alkyne cycloaddition (SPAAC) reaction with pendant azide (N3) groups on polymer carriers with high efficiency and without the use of a catalyst or reducing agent which complicates purification. The synthesis of the DBCO-terminated, cathepsin B-cleavable linker is detailed in **Scheme S2** and conjugation of the linker to diABZI-amine to generate the final diABZI-V/C-DBCO **(3)** product is described in **Scheme 1**. All compounds were characterized through nuclear magnetic resonance spectroscopy (NMR) and electrospray ionization mass spectrometry (ES-MS) **(Supplemental Figures S3-4).**

**Figure 2:**
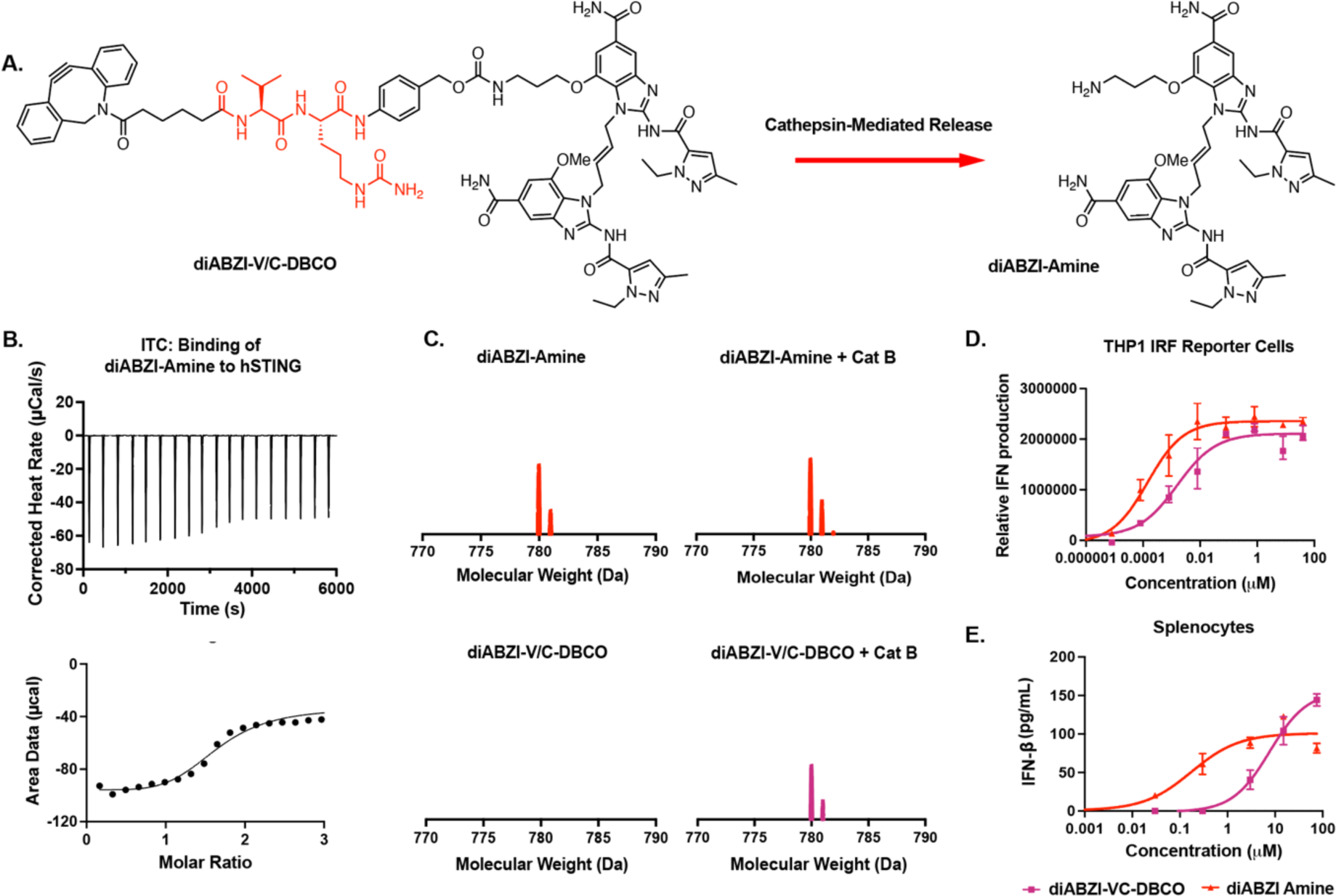
Design and characterization of diABZI-amine and diABZI-V/C-DBCO. **(A)** Reaction scheme of diABZI-amine release via cathepsin B-mediated cleavage of diABZI-V/C-DBCO. **(B)** Isothermal calorimetry (ITC) traces (top) and binding isotherm (bottom) of diABZI-amine binding to human recombinant STING. **(C)** MALDI-MS spectra (770-790 Da) of diABZI-amine and diABZI-V/C-DBCO with or without pre-incubation with cathepsin B to demonstrate enzyme sensitivity of the valine-citrulline-PAB linker used to allow for release diABZI-amine from diABZI-V/C-DBCO. **(D)** Dose-response curves for relative IFN-I production by THP1-Dual reporter cells treated with diABZI-amine and diABZI-V/C-DBCO (n=3). **(E)** Dose-response curves for IFN-β secretion by isolated murine splenocytes treated with diABZI-amine and diABZI-V/C-DBCO (n=2).

**Scheme 1:**
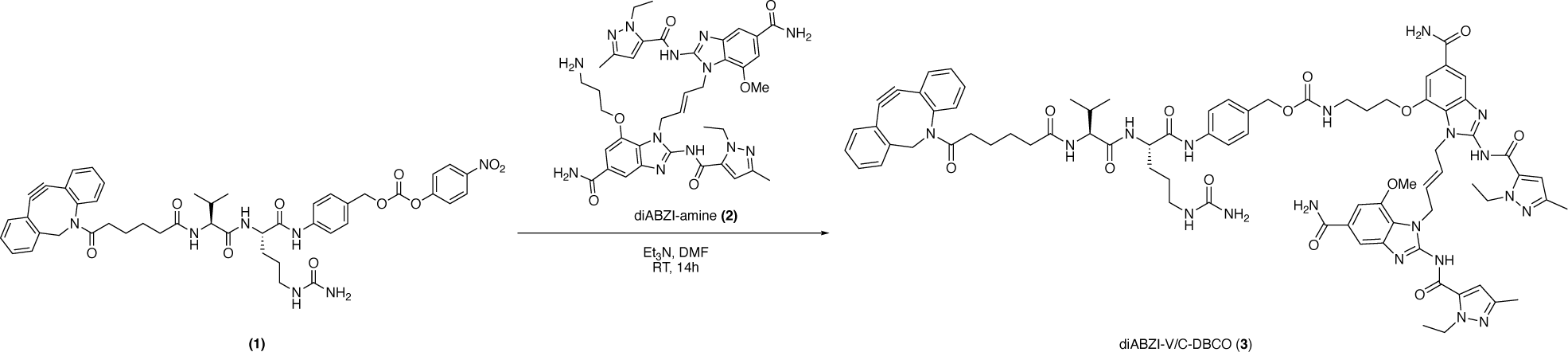
Synthesis of diABZI-V/C-DBCO.

Having synthesized a conjugatable diABZI, we first validated the ability of diABZI-amine to bind hSTING using isothermal calorimetry (ITC), measuring a KD of ∼70 nM (**Figure 2B**), similar to that previously described for diABZI compounds.^16^ We then confirmed cathepsin B-mediated release of diABZI-amine from diABZI-V/C-DBCO using MALDI to detect the emergence of peaks at 779 Da corresponding to the protonated form of diABZI-amine (**Figure 2C**). Finally, we characterized the activity of diABZI-amine and diABZI-V/C-DBCO compounds as STING agonists. We first utilized THP1-Dual reporter cells, a human monocyte cell line engineered with an IRF-inducible luciferase reporter that allows for relative measurement of IFN-I production. Both compounds were active in the nanomolar range with EC50 values of 0.144 ±0.149 nM for diABZI-amine and 1.47 ± 1.99 nM for diABZI-V/C-DBCO (**Figure 2D**). It is common for prodrugs to exhibit reduced *in vitro* activity compared to the parent drug due to their larger size and therefore slower cellular uptake and the kinetics of enzyme-mediated drug release. We also assessed activity in primary murine splenocytes using an IFN-β enzyme-linked immunosorbent assays (ELISA), measuring EC50 values for diABZI-amine and diABZI-V/C-DBCO of 0.17 ± 6.6 μM and 7.7 ± 0.05 μM, respectively (**Figure 2E**). Combined, these data demonstrate that diABZI-V/C-DBCO is a versatile and potent STING agonist for enzyme-cleavable conjugation to drug carriers via SPAAC click chemistry.

### Design and Characterization of STING-Activating Polymer-Drug Conjugates

Direct conjugation of drugs to water soluble polymers has been explored for several decades but only recently in the context of small molecule immunostimulatory agents.^35,36^ Some of the most well-defined and translationally-advanced polymer-drug conjugates utilize a 2-hydroxypropyl methacrylamide (HMPA) backbone, which has been established to be highly water-soluble and biocompatible.^37^ For example, HMPA copolymer-chemotherapy conjugates have demonstrated potential in clinical trials with minimal polymer toxicity.^26,38^ Polymer molecular weight directly affects the PK, biodistribution, and tumor accumulation with larger polymers avoiding renal clearance, resulting in increased circulation time and passive tumor accumulation.^25^ Since the renal filtration cut off for linear polymer chains is estimated to be ∼70kDa, we aimed to synthesize water-soluble polymers below (25kDa) and above (100kDa) this threshold to evaluate their utility as drug carriers for systemic administration of STING agonists. We chose N,N-dimethylacrylamide (DMA) as an inert and versatile polymer scaffold as it is a highly hydrophilic monomer akin to HPMA, but has a smaller side chain that reduces steric hindrance for drug conjugation. Poly(DMA) can also be efficiently synthesized via reversible-addition-fragmentation chain-transfer (RAFT) polymerization, allowing for precise control of molecular weight and the potential to incorporate a multitude of functional monomers into the backbone. To allow for conjugation of DBCO-functionalized STING agonists, we copolymerized an azide-functionalized monomer, azide-ethylmethacrylate (AzEMA), with DMA at 93:7 a DMA:AzEMA ratio using RAFT to synthesize poly(DMA93-*co*-AzEMA7) random copolymers of 25kDa and 100kDa (**Figure 3A**). Polymers were characterized through ^1^H NMR and gel permeation chromatography (GPC) (**Figure S5-8**), demonstrating precise control of molecular weight and monomer composition (**Table S9**), with the 25 kDa polymer containing ∼19 azide groups per chain and the 100 kDa polymer with ∼72 azide groups per chain. Both polymers were pre-labeled with ∼1 Cyanine5(Cy5)-DBCO fluorophore per chain through SPAAC to allow polymer and diABZI concentrations to be independently determined spectrophotometrically and for *in vitro* and *in vivo* evaluation of polymer uptake and distribution using fluorescence measurements. diABZI-V/C-DBCO agonists were covalently attached to both polymers via SPAAC at room temperature in dimethyl sulfoxide (DMSO) for 24 h to yield SAPCons with two distinct molecular weights (**Figure 3A**). A 7-molar excess of diABZI to the 25kDa polymer and 20 molar excess of diABZI to the 100kDa polymer was used to achieve approximately 1 mol% drug per chain. Conjugates were purified via dialysis against sterile water and diABZI and polymer concentrations were determined by UV-vis spectroscopy using absorbance peaks for diABZI at 325 nm and the Cy5-labeled polymer at 650 nm (**Figure 3B**). On average SAPCon[25kDa] had approximately 4.3 ± 1.2 drugs per chain and SAPCon[100kDa] had 7.3 ± 1.7 drugs per chain. This results in an average molecular weight of approximately 33,256 Da for SAPCon[25kDa] and 127,112 Da for SAPCon[100kDa]. An excess of azide groups was incorporated to allow for more efficient conjugation of DBCO-V/C-diABZI to polymer chains, though this is a variable that could be optimized in future iterations of SAPCon to minimize batch-to-batch variation. Dynamic light scattering indicated that conjugates exist as random, linear copolymers in solution and do not self-assemble into nanoparticles (**Figure S10**). This therefore enables direct investigation of backbone molecular weight on pharmacological properties, though future opportunities exist to optimize diABZI graft density to promote formation of self-assembled morphologies.

**Figure 3:**
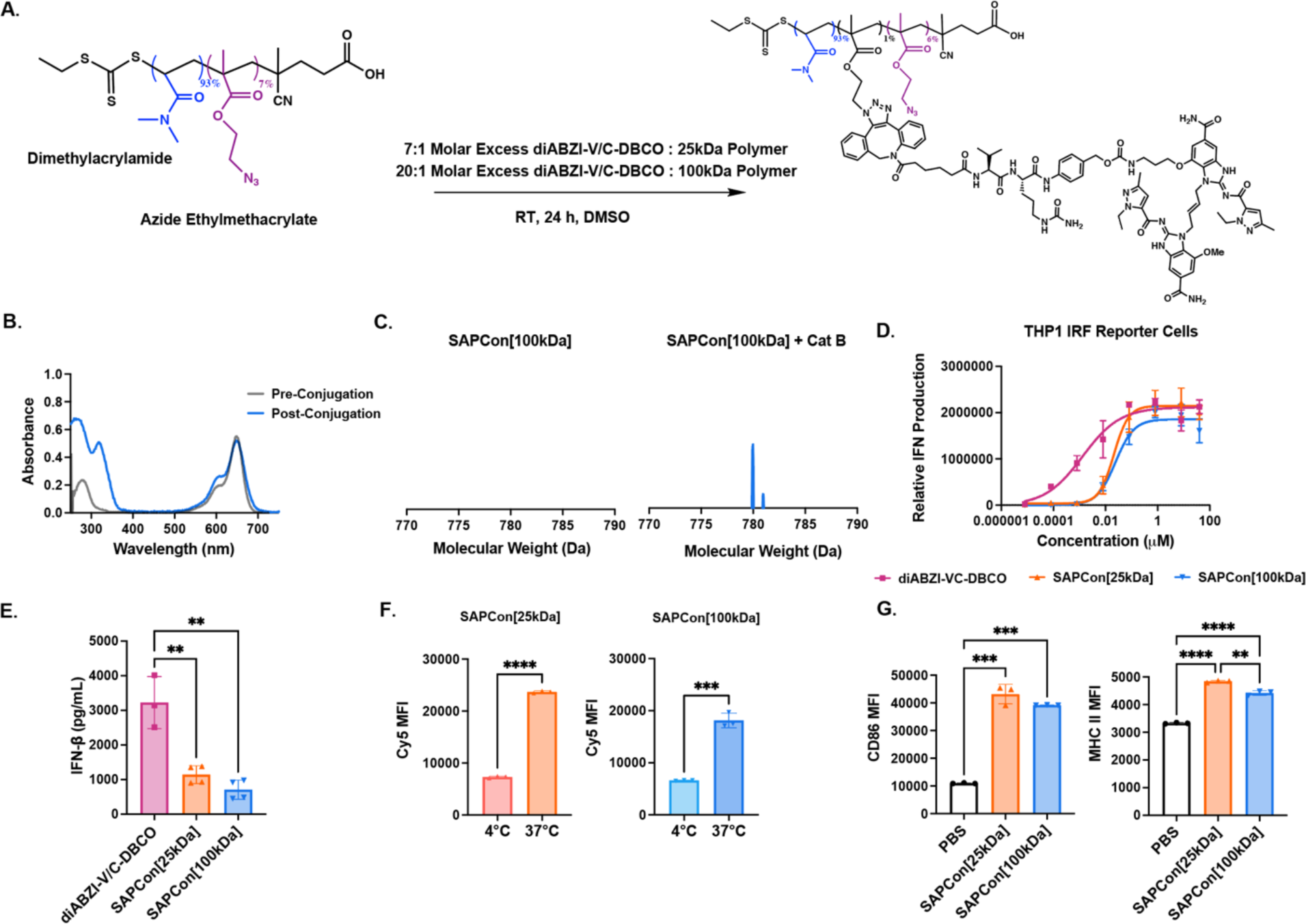
Design, synthesis, and characterization of STING-Activating Polymer-Drug Conjugates (SAPCon). **(A)** Reaction scheme for the synthesis of SAPCon[25kDa] and SAPCon[100kDa]. Poly(DMA_93_-*co*-AzEMA_7_) copolymers of 25 or 100 kDa molecular weight were reacted with DBCO-V/C-diABZI STING agonist to yield SAPCon. **(B)** UV-vis absorbance spectra of Cy5-labeled poly(DMA_93_-*co*-AzEMA_7_) copolymers pre-and post-conjugation to DBCO-V/C-diABZI demonstrating increased absorbance at 325 nm corresponding to diABZI. **(C)** MALDI-MS spectra (770-790 Da) of SAPCon[100kDa] with or without pre-incubation with cathepsin B to demonstrate enzyme-responsive release of diABZI-amine from SAPCon. **(D)** Dose-response curves for relative IFN-I production by THP1-Dual reporter cells treated with DBCO-V/C-diABZI, SAPCon[25kDa], and SAPCon[100kDa] (n=3; mean±SD). **(E)** IFN-β secretion by murine bone marrow-derived macrophages treated with DBCO-V/C-diABZI, SAPCon[25kDa], and SAPCon[100kDa] (n=3). **(F)** Median fluorescent intensity as measured by flow cytometry of BMDMs treated with Cy5-labeled SAPCon[25kDa] and SAPCon[100kDa] for 2 h at 37°C or 4°C (n=3). **(G)** Flow cytometric evaluation of CD86 and MCH-II expression by BMDMs treated with SAPCon[25kDa], SAPCon[100kDa], or PBS (vehicle) (n=3).

Since grafting to polymer chains can inhibit enzyme access to cleavable spacers, we next evaluated cathepsin B-mediated release of conjugated diABZI from SAPCon[100kDa] by using MALDI to confirm the liberation of diABZI-amine as indicated by peaks corresponding to 779 Da. (**Figure 3C**). We next evaluated the activity of SAPCons *in vitro* using the previously described THP1-Dual IRF reporter cells. SAPCon[25kDa] and SAPCon[100kDa] were both active with EC50 values of 20.8 ± 13.3 nM and 24.1 ± 12.9 nM (**Figure 3D**), respectively, compared to 1.3 ± 9 nM for diABZI-V/C-DBCO. We also assessed activity of SAPCons in bone marrow-derived macrophages (BMDMs), finding both variants to induce production of IFN-β (**Figure 3E**), a signature cytokine of STING activation. It is unsurprising that both conjugates exhibited decreased activity when compared to the unconjugated diABZI-V/C-DBCO as it is common for polymer-drug conjugates to have reduced *in vitro* activity due to lack of membrane permeability and dependence on endocytosis and drug release. Indeed, we observed a significant decrease in Cy5-labeled SAPCs in BMDMs at 4°C as measured by flow cytometry, indicating the necessity for endocytosis (**Figure 3F**). Since macrophage activation is one of the hallmarks of STING activation ^39^, we also tested the ability of the conjugates to promote BMDM activation *in vitro* via flow cytometric analysis CD86 and MCH-II expression, finding that both conjugates exhibited a significant upregulation of these proinflammatory macrophage markers (**Figure 3G**).

### Evaluation of SAPCon Pharmacokinetics, Biodistribution, and Cellular Uptake

Delivered without a carrier, STING agonists are cleared relatively quickly upon systemic administration with minimal accumulation at tumor sites, resulting in insufficient STING activation and/or the potential for unintended side effects.^40^ Conjugation of drugs to hydrophilic polymeric carriers can increase their circulation time and promote accumulation at tumor sites in a molecular weight-dependent manner.^41^ We therefore evaluated the pharmacokinetics and biodistribution of Cy5-labeled SAPCon. To evaluate pharmacokinetics, healthy mice were intravenously injected with either SAPCon[25kDa] or SAPCon[100kDa] at a diABZI dose of approximately 0.012 μmol/mouse, and blood was sampled over 24 h for spectrofluorometric quantification of polymer concentration as a function of time. As expected, SAPCon[100kDa] demonstrated prolonged circulation, with a serum elimination half-life of approximately 4.4 h compared to 1.1 h for SAPCon[25kDa] as determined using a two-phase decay model (**Figure 4A**).

**Figure 4:**
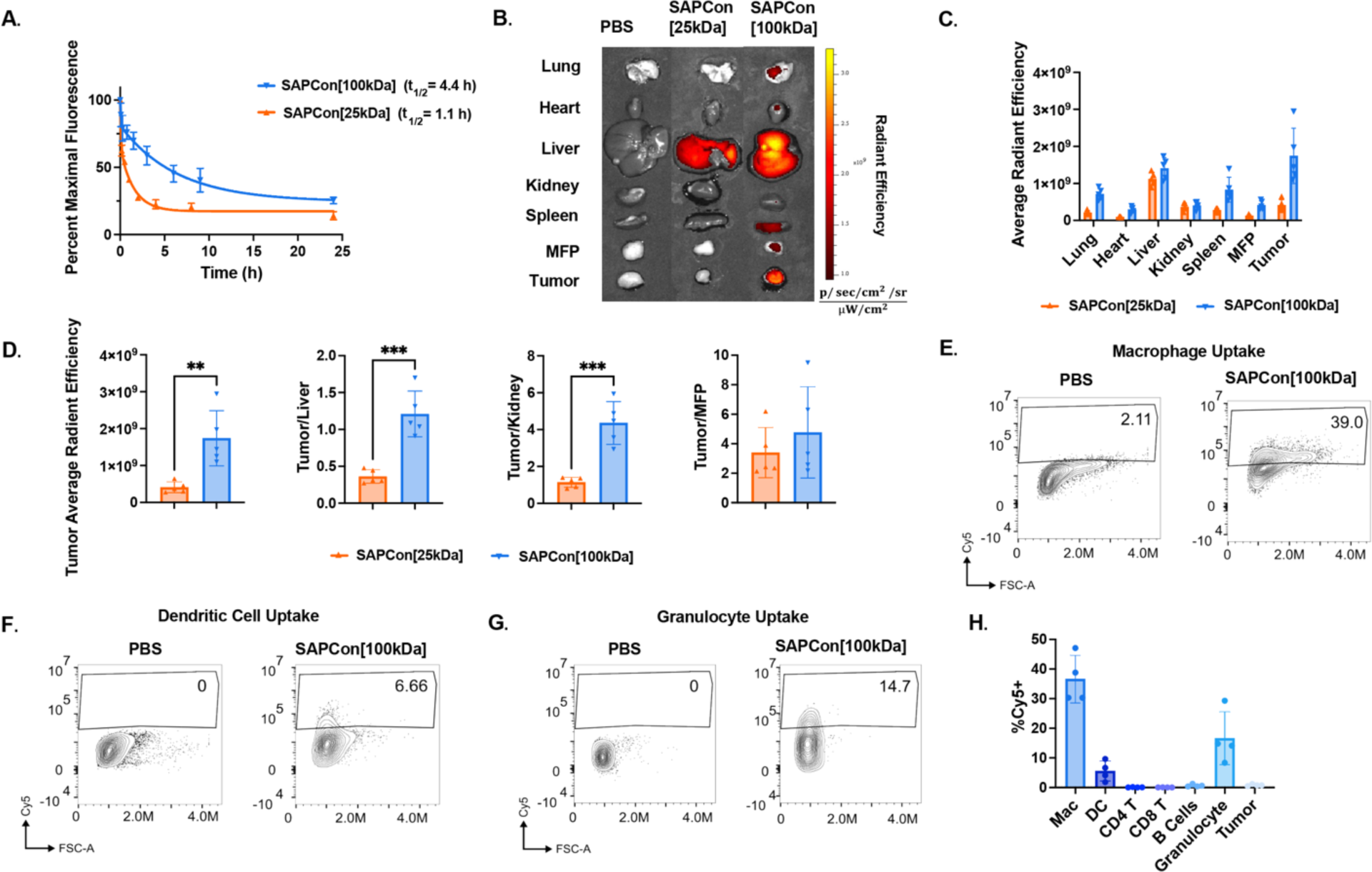
SAPCon promotes tumor accumulation and cellular uptake by tumor-associated myeloid cells. **(A)** Plasma pharmacokinetics of intravenously-administered Cy5-labeled SAPCon[25kDa] and SAPCon[100kDa] in healthy C57/BL6 mice (n=5). Data were fit to a two-phase decay model to determine an elimination half-life of 1.1 h for SAPCon[25kDa] and 4.4 h for SAPCon[100kDa]. **(B)** Representative IVIS fluorescent images of excised EO771 tumors and organs 48 h following administration of Cy5-labeled SAPCon. **(C)** Quantification of tissue fluorescence measured with IVIS imaging 48 h following administration of Cy5-labeled SAPCon (n=5). **(D)** Ratios of tissue fluorescence between tumor and/or organs. **(E-G)** Representative flow cytometry plots of Cy5^+^ cells within gates corresponding to **E)** CD11b^+^F4/80^+^ macrophages, **F)** CD11b^+^ dendritic cells, and **G)** CD11b^+^Gr1^+^ granulocytes. **H)**. Summary of cellular uptake of Cy5-labeled SAPCon[100kDa] by EO771 tumor-associated cell populations using flow cytometry (n=4).

A biodistribution study was next performed to evaluate SAPCon accumulation in tumor tissue and major organs after 24 and 48 hours. We selected breast cancer as a model for our studies because anti-PD-1 (e.g., pembrolizumab) is currently approved for treatment of TNBC but with a response rate of only 5-20%, motivating the need for adjunctive or alternative therapies to improve responses.^42^ Female C57BL/6 mice were inoculated with EO771 breast cancer cells in the left-side 4th mammary fat pad, tumors were grown to approximately 50 mm^3^, and mice were intravenously administered SAPCon[25kDa] or SAPCon[100kDa] at approximately 0.012 μmol diABZI/mouse; PBS was used a vehicle control. Tumor and organs were resected after 24 or 48 hours for IVIS imaging (**Figure 4B, Figure S11A**). As anticipated, renal clearance was evident with SAPCon[25kDa] at 24 hours, with higher Cy5 ratios in the kidneys compared to SAPCon[100kDa] (**Figure S11B-C**). After 48 hours, significant tumor accumulation was measured within the tumor for SAPCon[100kDa] compared to SAPCon[25kDa] **(Figure 4C-D)**. Unsurprisingly, both conjugates accumulated in the liver, although preferential tumor accumulation was measured for SAPCon[100kDa] **(Figure 4D**). Both carriers accumulated ∼4-fold in tumor tissue over healthy contralateral mammary fat pad **(Figure 4D**). We then completed a dose-finding study to determine relative toxicity of SAPCon[25kDa] and SAPCon[100kDa] through weight loss measurements and found that SAPCon[25kDa] corresponded resulted in more weight loss than SAPCon[100kDa] (**Figure S12**). A dose correlating to 8 μg of diABZI (0.009μg, or 0.4 mg/kg) was selected for the following studies since this dose and regimen were well tolerated with mice exhibiting mild and transient weight loss in the immediate post-treatment period, a pattern that has been commonly observed with many promising STING agonists delivery systems. Due to the longer circulation time, increased tumor accumulation, and lower transient weight loss of the 100kDa conjugate, SAPCon[100kDa] was selected as the lead platform for the remaining studies.

We next aimed to determine which cell types within the TME internalize SAPCon[100kDa] using flow cytometry. Mice with EO771 breast tumors were intravenously administered Cy5-labeled SAPCon[100kDa] at a dose corresponding to approximately 0.009 μmol diABZI/mouse (0.4 mg/kg), and tumors were harvested 24 hours post-injection. Surface makers were used to identify major immune cell populations (CD45^+^), including macrophages (CD11b+F4/80^+^), dendritic cells (CD11c^+^), granulocytes (CD11b^+^Gr1^+^), CD4^+^ T cells, CD8^+^ T cells, and B cells (CD19); CD45^-^ cells are primarily breast cancer cells but may also include fibroblasts, endothelial cells, and other stromal cells. (**Figure S13**). Interestingly, conjugates were primarily internalized by tumor-associated macrophages (TAMs), with >25% of Cy5^+^ macrophages, consistent for their propensity to endocytose nanoparticles and large macromolecules in the TME (**Figure 4E,H**). TAMs are often abundant in tumors and can suppress antitumor immune responses, and therefore such passive targeting of STING agonists to TAMs has the potential to promote polarization to an antitumor, proinflammatory phenotype.^43^ We also observed polymer-diABZI uptake by DCs, professional antigen presenting cells that can prime antitumor T cell responses, as well as granulocytes that may also promote antitumor immunity in response to STING activation (**Figure 4F-H, S14**).^44,45,46^ Consistent with their relatively poor capacity for endocytosis, we did not observe uptake by T cells; this may reflect an additional advantage of SAPCon over free diABZI since T cells are susceptible to STING-induced apoptosis. Surprisingly, we did not observe significant conjugate uptake in CD45^-^ cells, which are predominantly breast cancer cells. This is potentially significant since STING is known to be silenced or dysfunctional in some human cancers which poses a challenge for STING agonist delivery technologies that target cancer cells.

STING, along with other innate immune pathways, occupies a distinctive niche in the drug delivery field due to the challenge of striking the right balance in the magnitude and kinetics of the response as hyperacute STING activation can promote cell death and increase toxicity while sustained activation can lead to inflammatory-driven diseases and downregulation of the pathway.^47,14^ However, the optimal PK profile for STING agonists is not known and systematic studies exploring the effect of half-life on efficacy and safety are lacking. Therefore, SAPCon may also offer a versatile tool for precisely modulating drug half-life through control of polymer properties to optimally balance safety and efficacy. We also sought to minimize complexity in our initial SAPCon design through use of a linear poly(DMA) scaffold that harnesses increased circulation time to passively accumulate at tumor sites. However, SAPCon is versatile platform from which to further build and optimize; notably, our approach is highly amenable to integration of small molecule targeting ligands such as mannose, folate, or integrin ligands (e.g., RGD) as well a different linkers (e.g., ROS-responsive, MMP-cleavable) that may further increase tumor accumulation or delivery to specific cell populations.

### SAPCon Enriches STING Activation in the TME and Stimulate an Immunogenic Microenvironment

Small molecule STING agonists have shown promise in murine tumor models, but they lack the ability to preferentially accumulate within the tumor, motivating our development STING-activating polymer-drug conjugates that can enrich STING activation in the TME. We therefore assessed the capacity of SAPCon[100kDa] to increase STING activation in the TME relative to small molecule diABZI agonists by measuring STING-driven gene expression in breast tumors when diABZI-V/C-DBCO was systemically administered with and without the polymeric carrier. C57BL/6 mice with orthotopic EO771 tumors (∼50-100 mm^3^) were systemically administered diABZI-V/C-DBCO or diABZI-polymer conjugate at a matched dose of 0.009 μmol diABZI/mouse (0.4 mg/kg), and tumors were harvested 6 h post injection. Using RT-qPCR to evaluate STING-related gene expression, we found that *Tmem173, Ifnb, Cxcl10, Tnf, and Il6* were all significantly upregulated in the tumor for mice treated with the conjugate (**Figure 5A-B**). By contrast, no significant increase was found in in the tumors of mice with treated with diABZI-V/C-DBCO relative to PBS, demonstrating the ability of polymer-diABZI conjugates to enhance STING activation in tumor tissue. We also collected blood at 6 h post-treatment for serum cytokine analysis, finding that SAPCon[100kDa] also increased serum IFN-β levels relative to diABZI-V/C-DBCO, potentially due to increased circulation time (**Figure 5C**). Interestingly, while both diABZI-V/C-DBCO and SAPCon[100kDa] increased serum IFN-β levels, only SAPCon[100kDa] increased STING activation in the TME.

**Figure 5:**
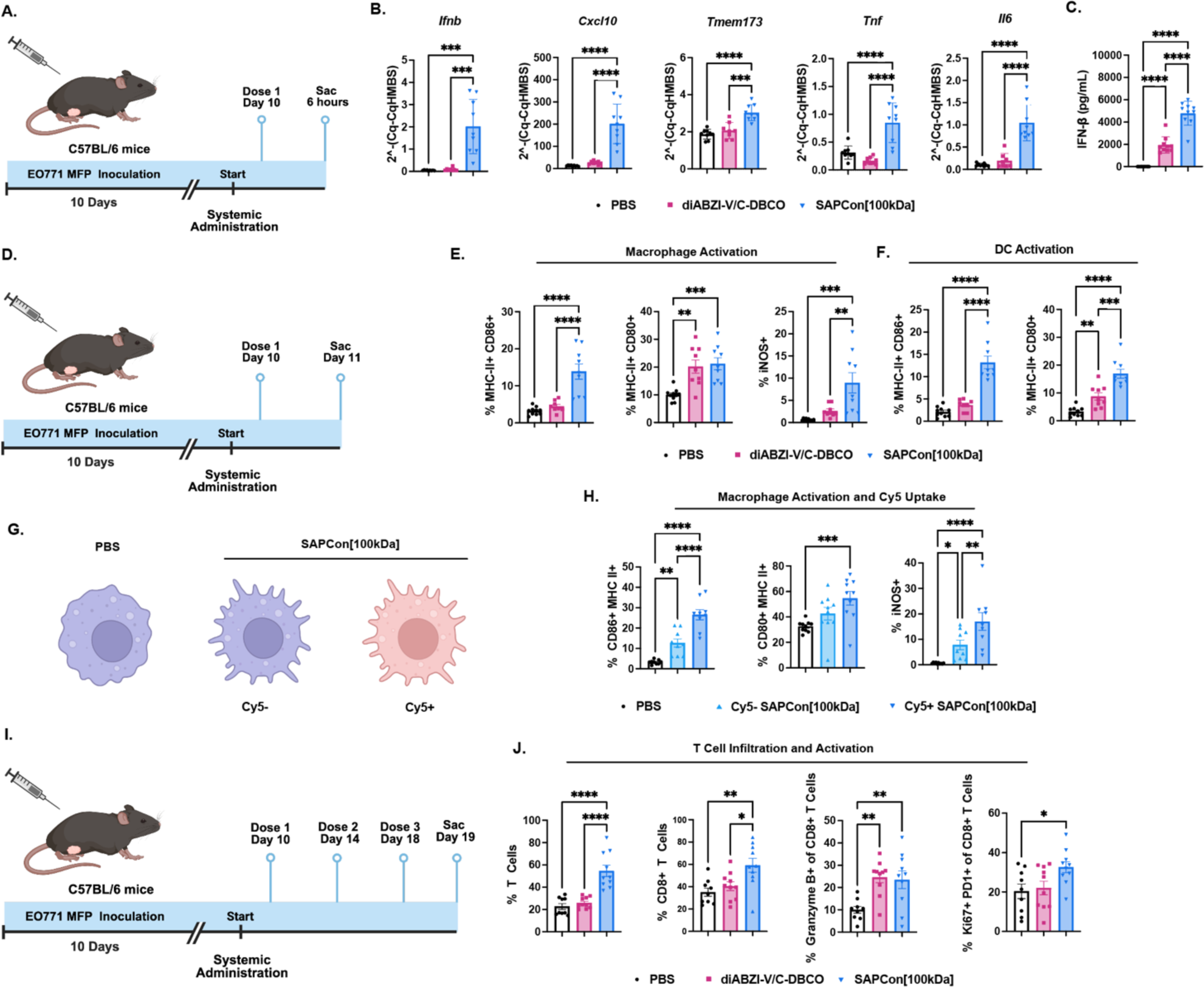
SAPCon enhances STING activation at tumor sites and reprograms the tumor microenvironment. **(A)** Schematic of EO771 tumor inoculation, treatment schedule, and study end point for analysis. **(B)** Gene expression analysis of tumor tissue (*Ifnb1*, *Cxcl10*, *Tmem173*, *Tnf*, and *Il6*) measured by qRT-PCR and **(C)** Serum IFN-β concentration at 6 h post-intravenous administration of SAPCon[100kDa], diABZI-V/C-DBCO, or PBS (n=10). **(D)** Schematic of EO771 tumor inoculation, treatment schedule, and study end point for flow cytometric immunophenotyping. Flow cytometric analysis of the frequency of **(E)** MHC-II^+^CD86^+^, MHC-II^+^CD80^+^, and iNOS^+^ macrophages and **(F)** MHC-II^+^CD86^+^ and MHC-II^+^CD80^+^ dendritic cells in EO771 breast tumors 24 h following a single administration of SAPCon[100kDa], diABZI-V/C-DBCO, or PBS (n=10). **(G)** Schematic of non-activated macrophages from control mice vs activated macrophages from mice treated with SAPCon[100kDa]. Treatment can not only activate macrophages that endocytose SAP[Con100kDa], but also elicit a bystander effect. **(H)** Flow cytometric analysis of the frequency MHC-II^+^CD86^+^, MHC-II^+^CD80^+^, and iNOS^+^ cells within Cy5^+^ and Cy5^-^ macrophage populations following administration of Cy5-labeled SAPCon[100kDa] relative to macrophages from mice treated with PBS. **(I)** Schematic of EO771 tumor inoculation, treatment schedule, and study end point for flow cytometric immunophenotyping. **(J)** Flow cytometric analysis of the frequency CD3^+^ T cells, CD8^+^ T cells, granzyme B^+^CD8^+^ T cells, and Ki67^+^PD-1^+^CD8^+^ T cells in EO771 tumors 24 h following the last of three doses of SAPCon[100kDa], diABZI-V/C-DBCO, or PBS (n=10). All data were plotted using GraphPad 10 with multiple comparisons through one-way ANOVA.

Based on its capacity to increase STING activation in tumors, we next sought to understand how SAPCon[100kDa] impacts the immunocellular composition of the breast TME. Mice with orthotopic EO771 tumors (∼50-100 mm^3^) were treated with either diABZI-V/C-DBCO or the conjugate at matched a drug concentration of 0.009 μmol/mouse (0.4 mg/kg). First, given the propensity of SPACon[100kDa] to be internalized by tumor associated myeloid cells, tumors were resected 24 h after treatment to analyze myeloid cell activation via flow cytometry (**Figure 5D-F, S15**). SPACon[100kDa], but not diABZI-V/C-DBCO, resulted in increased frequency of CD86^+^MHCII^+^ and iNOS^+^ CD11b^+^F4/80^+^ macrophages, while both treatments induced a similar frequency of CD80^+^MHCII^+^ macrophages relative to vehicle control. Interestingly, we found that a higher percentage of activated macrophages were also Cy5^+^, indicating that conjugates were more effective in cells that internalized them; however, the level of Cy5^-^ activated macrophages was higher than in vehicle treated mice, suggesting a bystander effect mediated by local cytokines and/or delivery of liberated diABZI to surrounding cells (**Figure 5G-H, S16**). We also found that SAPCon[100kDa] increased DC activation relative to diABZI-V/C-DBCO as evidenced by elevated levels of the costimulatory molecules CD86 and CD80 (**Figure 5F, S15**). Hence, SAPCon[100kDa] generates proinflammatory APCs which can play important roles in tumor antigen processing and presentation, T cell priming, cancer cell lysis, and inhibition of cancer cell proliferation.

Although the cancer-immunity cycle involves a myriad of immune cells that coordinate an antitumor response, cytotoxic CD8^+^ T cells are considered the main effectors for cancer cell killing and primary targets for anti-PD-1 ICB.^48^ To evaluate the ability of SAPCon[100kDa] to potentiate CD8^+^ T cell response in the tumor, mice with EO771 breast tumors (∼50 mm^3^) were treated with either diABZI-V/C-DBCO or SAPCon[100kDa] at matched a drug concentration of 0.009 μmol/mouse (0.4 mg/kg) for three total doses, four days apart (**Figure 5I**), which is a common treatment regimen for preclinical evaluation of STING agonists.^49^ Tumors were resected 24 h after the last injection and analyzed via flow cytometry. Treatment with SAPCon[100kDa] resulted in a significant increase in both total CD3^+^ T cell and CD8^+^ T cell frequency within the tumor compared to diABZI-V/C-DBCO or vehicle control. Within the CD8^+^ T cell population, we found that SPACon[100kDa] slightly increased the frequency of Ki67^+^PD-1^+^CD8^+^ T cells, which have been previously correlated with improved responses to ICB^50^, and both treatments increased the frequency of granzyme B^+^ CD8^+^ T cells (**Figure 5J, S17**). Collectively, these data demonstrate that this conjugate can increase STING activation, primarily by targeting tumor-associated myeloid cells, resulting in activation of macrophages and DCs and increased infiltration of cytotoxic T cells.

### SAPCon Enhances Antitumor Efficacy and Improve Survival in Murine Breast Cancer Models and Improves Responses to Anti-PD-1 ICB

Based on the capacity of polymer-diABZI conjugates to accumulate and activate STING within the tumor and to invigorate an antitumor immunological shift of the TME, we next aimed to determine if this translated to improved therapeutic efficacy and survival. We first evaluated the tolerability and safety of SAPCon[100kDa] by administering healthy mice SAPCon[100kDa] at 0.4 mg/kg diABZI using a three-dose regimen with doses spaced three days apart and euthanizing mice a week following the final administration for evaluation of blood chemistry and organ pathology. There were no significant differences in common blood toxicity markers between treatment and control groups, including ALT, AST, and bilirubin, which are markers of liver toxicity where potential on-target, off-tumor toxicity might be anticipated based on the liver accumulation (**Figure S18**). Histopathology analysis of the liver showed no evidence of hepatotoxicity with some extramedullary hematopoiesis in both liver and spleen, common for treatments via innate immune agonists. However, analysis of the bone marrow indicated no significant differences between treated and untreated groups. Hence, SAPCon[100kDa] was well-tolerated at this dose, which we selected for initial evaluation of therapeutic efficacy and set as our maximum dose to avoid the risk of additional toxicities.

Using this dose and regimen, C57BL/6 mice with EO771 mammary tumors (∼50 mm^3^) were treated with either free diABZI-V/C-DBCO or SAPCon[100kDa]. Tumor volume was monitored over time, with a humane endpoint at 1500 mm^3^ (**Figure 6A-D**). Weight loss was also measured throughout treatment (**Figure S19**). We found that treatment with SAPCon[100kDa] resulted in a significant reduction in tumor size and prolonged survival with a 37.5% complete response rate observed, whereas diABZI-V/C-DBCO had no effect relative to PBS control. We note that this represents a significantly improved response in the EO771 model relative to a previously described nanoparticle-based delivery system.^15^ Mice exhibiting complete responses were rechallenged with EO771 cells on the contralateral mammary fat pad and all were resistant to tumor rechallenge, demonstrating the establishment of immunological memory in response to treatment with SAPCon[100kDa] (**Figure 6E**).

**Figure 6:**
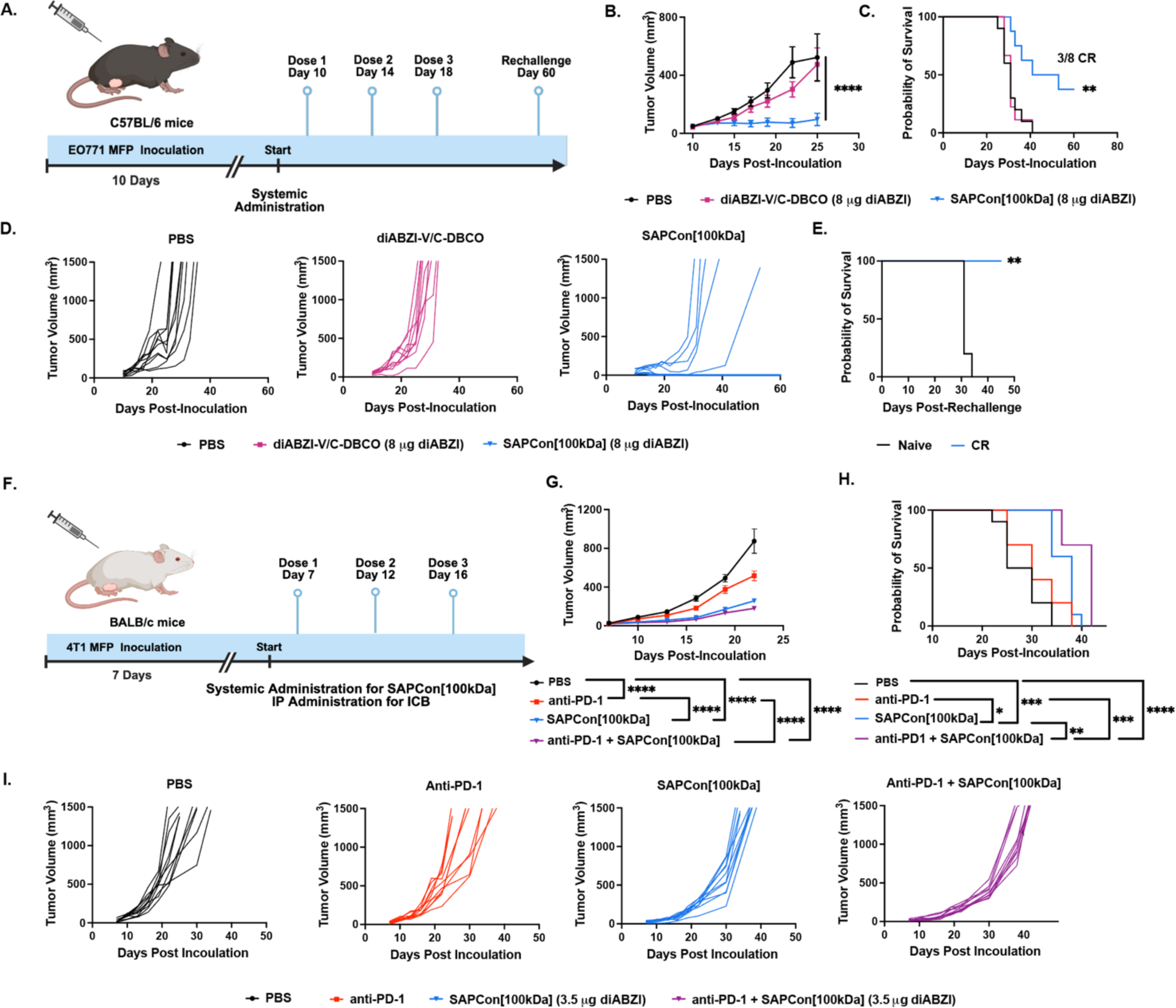
Systemically administered SAPCon inhibits tumor growth and improves response to anti-PD-1 immune checkpoint blockade in models of breast cancer. **(A)** Schematic of EO771 tumor inoculation and treatment schedule. **(B)** Average tumor growth curves. **(C)** Kaplan-Meier survival plots for mice with EO771 tumors treated as indicated. CR = complete responder **(D)** Spider plots of individual tumor growth curves (PBS: n=10, diABZI-V/C-DBCO: n=9, SAPCon[100kDa]:n=8). **(E)** Kaplan-Meier survival plots for tumor rechallenge model (Naïve: n=5, CR: n=3). (F) Schematic of 4T1 tumor inoculation and treatment schedule. **(G)** Average tumor growth curves. **(H)** Kaplan-Meier survival plots for mice with 4T1 tumors treated as indicated. **(I)** Spider plots of individual tumor growth curves (n=10).

We next sought to evaluate if SAPCon[100kDa] could improve response to anti-PD-1 ICB therapy, which is approved for use in TNBC but with a response rate of only ∼5-20%.^51^ For these studies, we selected a 4T1 model, an aggressive and poorly immunogenic model of TNBC. We also reduced the dose of SAPCon[100kDa] to 3.5μg per mouse (0.004 μmol, 0.175 mg/kg) to mitigate the risk of any additional toxicity that might be occur in response to combination therapy. Female BALB/c mice were inoculated with 4T1 mammary fat pad tumors which were treated with either PBS, anti-PD-1, SAPCon[100kDa], or combination of anti-PD1 and SAPCon[100kDa] once tumors reached approximately 20 mm^3^. Treatment was administered every 4 days for 3 total doses. Tumor volume was monitored over time, with a humane endpoint at 1500mm^3^ (**Figure 6F-I**). Weight loss was also monitored through treatment (**Figure S20**). Both SAPCon[100kDa] and combination therapy significantly reduced tumor burden and prolonged survival, with combination therapy eliciting the highest efficacy. Although anti-PD-1 resulted in a slight reduction in tumor growth, combination with SAPCon significantly improved this current standard-of-care immunotherapy for treatment of TNBC, further demonstrating the potential of SAPCon as a next-generation cancer immunotherapeutic.

## Conclusions

The stimulator of interferon genes (STING) pathway is as an important mediator in antitumor immunity, but pharmacological barriers continue to limit the clinical realization of STING agonists and motivate the need to innovate drug delivery systems for this promising class of immunotherapy agents. To address current challenges, we developed STING-activating polymer-drug conjugates (SAPCon), a platform technology for systemic administration of diABZI-based STING agonists. Here we described the synthesis, characterization, and biological evaluation of a first-generation SAPCon fabricated via SPAAC click chemistry between pendant azide moieties copolymerized into a hydrophilic poly(DMA) backbone and a novel DBCO-functionalized, cathepsin B-responsive diABZI STING agonist. Selecting a 100kDa polymer backbone to avoid renal clearance and achieve extended half-life, we demonstrate that systemically administered SAPCon[100kDa] accumulates in tumor tissue to potentiate STING activation in the tumor microenvironment (TME), primarily via endocytosis by tumor-associated macrophages. Consequently, SAPCon[100kDa] stimulated macrophage and dendritic cell activation in breast tumors and increased the infiltration of activated CD8^+^ T cells. This shift in the immune contexture of the TME ultimately resulted in robust antitumor efficacy in an EO771 breast cancer model as evidenced by inhibition of tumor growth and increased survival benefit, with nearly 40% of mice exhibiting complete responses and durable immunological memory that protected against tumor rechallenge. We also found that SAPCon[100kDa] was effective in a challenging 4T1 model of triple negative breast cancer in combination with currently approved anti-PD-1 ICB. Finally, while we leveraged a poly(DMA) carrier and cathepsin B-cleavable linker, the synthetic workflow employed allows for future exploration of SAPCon comprised of different polymer backbones, polymer architectures, and drug linker chemistries to improve the efficacy and/or safety of systemically-administered STING agonists for cancer immunotherapy.

## Methods

**Synthesis of diABZI-amine (2):** tert-butyl (E)-(3-(5-carbamoyl-2-((4-((4-carbamoyl-2-methoxy-6-nitrophenyl)amino)but-2-en-1-yl)amino)-3-nitrophenoxy)propyl)carbamate: In a sealable tube mixture of tert-butyl (3-(5-carbamoyl-2-chloro-3-nitrophenoxy)propyl)carbamate (**4**) (14.2 g, 38.0 mmol, 1.1 eq), (E)-4-((4-aminobut-2-en-1-yl)amino)-3-methoxy-5-nitrobenzamide, hydrochloride (**5**) (11.0 g, 34.6 mmol, 1 eq), hunig’s base (30 mL, 172.8 mmol, 5eq), and n-butanol (140 mL) was maintained at rt and argon was bubbled through the solution for 5 min. The mixture was sealed under an inert atmosphere, and heated to 120 °C for 24h, during which time the reaction mixture become homogeneous solution with brick red color. After cooling to rt the solid product was isolated by filtration washed with 25 mL ethanol followed by washing with 150 mL ether and dried to afford the desired product **(6)** (16.5 g, 26.7 mmol, 77%) as a brick red solid. ^1^H NMR (400 MHz, DMSO) δ 8.15 (s, 2H), 8.00 (broad s, 2H), 7.72 – 7.68 (m, 2H), 7.51 (s, 2H), 7.31 (broad s, 2H), 6.91 (t, *J* = 5.0 Hz, 1H), 5.68 – 5.58 (m, 2H), 4.12 – 4.04 (m, 4H), 4.00 (t, *J* = 6.2 Hz, 2H), 3.82 (s, 3H), 3.08 (dt, *J* = 6.2, 5.0 Hz, 2H), 1.85 (p, *J* = 6.2 Hz, 2H), 1.35 (s, 9H). HRMS (ESMS) Calculated for C_27_H_35_N_7_O_10_ [M+H]^+^: 618.2523, found 618.2507.

A solution of *tert*-butyl (*E*)-(3-(5-carbamoyl-2-((4-((4-carbamoyl-2-methoxy-6-nitrophenyl)amino)but-2-en-1-yl)amino)-3-nitrophenoxy)propyl)carbamate (**6)** (16.5 g, 26.7 mmol, 1eq) in 300 mL methanol was maintained at rt. To which aqueous solution of sodium hydrosulphite (65 g, 374 mmol, 14 eq) in 150 mL water added. After 5 min ammonium hydroxide (87 mL, 668 mmol, 25 eq) added to the reaction mixture. The reaction mixture was allowed to stir for 45 min during which time the color of the reaction mixture changed from orange to light yellow. After consumption of starting material, as judged by TLC analysis, the reaction mixture was filtered through celite. NaCl added to saturate the aqueous layer which was extracted by EtOAc (3 x 200 mL). The organic layer was dried over Na_2_SO_4_ and evaporated *in vacuo* to obtain the crude product. The crude reaction mixture was purified by column chromatography on basic alumina (5% to 35% MeOH:DCM) to get the desired product (**7)** as a thick yellow oil (6.7g, 12 mmol, 45%). (R_f_ = 0.4 in 10% MeOH in DCM on silica gel TLC). ^1^H NMR (400 MHz, DMSO) δ 7.60 (broad s, 2H), 6.96 (broad s, 2H), 6.89 (t, *J* = 5.1 Hz), 6.85 (dd, *J* = 4.0, 1.7 Hz, 2H), 6.77 (merged dd, 2H), 5.72 – 5.63 (m, 2H), 4.64 (d, *J* = 4 Hz, 4H), 3.94 (t, J = 6.1 Hz, 2H), 3.82 – 3.75 (m, 2H), 3.74 (s, 3H), 3.55 – 3.45 (m, 4H), 3.10 (dt, *J* = 6.1, 5.1 Hz, 2H), 1.83 (p, *J* = 6.2 Hz, 2H), 1.36 (s, 9H). HRMS (ESMS) Calculated for C_27_H_39_N_7_O_6_ [M+H]^+^: 558.3040, found 558.3023.

To a stirred solution of *tert*-butyl (*E*)-(3-(3-amino-2-((4-((2-amino-4-carbamoyl-6-methoxyphenyl)amino) but-2-en-1-yl)amino)-5-carbamoylphenoxy)propyl)carbamate **(7)** (6.7g, 12.0 mmol, 1eq) in DMF (60 mL) add solution of 1-ethyl-3-methyl-1*H*-pyrazole-5-carbonyl isothiocyanate (**8)** (5.2 g, 26.4 mmol, 2.2 eq) in DMF (12 mL) under inert atmosphere and stir for 45 min. EDC (5.8 g, 30.0 mmol, 2.5 eq) followed by triethylamine (8.4 mL, 60.0 mmol, 5 eq) were added to the reaction mixture and stirred overnight. The reaction mixture was diluted with diethyl ether to precipitate out the crude reaction mixture. Solid was filtered and resuspended in aqueous ammonium chloride solution (10 g NH_4_Cl in 100 mL water) stirred for 15 min and filtered followed by washing with water (2 × 50 mL), diethyl ether (2 × 50 mL) to obtain the desired intermediate (8.1 g, 9.22 mmol, 77%) as an off white solid. ^1^H NMR (400 MHz, DMSO) δ 7.95 (broad s, 2H), 7.95 (broad s, 2H), 7.63 (d, *J* = 2.63 Hz, 2H), 7.33 (broad s, 2H), 7.30 (d, *J* = 4.46 Hz, 2H), 6.85 (t, *J* = 5.1 Hz, 1H), 6.5 (s, 2H), 5.89 – 5.77 (m, 2H), 4.95 – 4.89 (m, 4H), 4.51 (q, *J* = 6.89 Hz, 4H), 3.99 (t, *J* = 6.1 Hz, 2H), 3.72 (s, 3H), 3.61 (dt, *J* = 6.1, 5.1 Hz, 2H), 2.09 (s, 6H), 1.70 (p, *J* = 6.1 Hz, 2H), 1.32 (s, 9H), 1.26 (t, *J* = 6.89 Hz, 6H). HRMS (ESMS) Calculated for C_43_H_53_N_13_O_8_ [M+H]^+^: 880.4218, found 880.4174.

To a stirred suspension of intermediate *tert*-butyl (*E*)-(3-((5-carbamoyl-1-(4-(5-carbamoyl-2-(1-ethyl-3-methyl-1*H*-pyrazole-5-carboxamido)-7-methoxy-1*H*-benzo[*d*]imidazol-1-yl)but-2-en-1-yl)-2-(1-ethyl-3-methyl-1*H*-pyrazole-5-carboxamido)-1*H*-benzo[*d*]imidazol-7-yl)oxy)propyl)carbamate (6.6 g, 7.5 mmol, 1 eq) in dichloromethane (70 mL) under inert atmosphere at room temperature add trifluoroacetic acid (5.7 mL, 75 mmol, 10 eq) dropwise which convert suspension in to true solution and stir for 6 hr. The diethyl ether added to precipitate out the solid product as tri TFA salt. The solid was filtered over Büchner funnel and washed with diethyl ether (2 x 100 mL) to obtain the tri TFA salt of desired compound, diABZI-amine, (**2)** as an off-white solid (6.5 g, 5.8 mmol, 77%). ^1^H NMR (400 MHz, DMSO) δ 7.96 (broad s, 2H), 7.71 (broad s, 2H), 7.64 (d, *J* = 5.4 Hz, 2H), 7.36 (broad s, 2H), 7.32 (dd, *J* = 9.5, 0.9 Hz, 2H), 6.5 (d, *J* = 9.0 Hz, 2H), 5.84 – 5.73 (m, 2H), 4.92 – 4.89 (m, 4H), 4.54 – 4.47 (m, 4H), 4.09 (t, *J* = 6.1 Hz, 2H), 3.71 (s, 3H), 2.92 – 2.84 (m, 2H), 2.11 (s, 3H), 2.09 (s, 3H), 1.89 (p, *J* = 6.1 Hz, 2H), 1.25 (dt, *J* = 7.08, 4.6 Hz, 6H). ^13^C NMR (151 MHz, DMSO) δ 168.1, 158.9, 158.7, 158.5, 158.3, 145.6, 145.3, 144.4, 130.5, 130.5, 128.4, 128.2, 109.7, 109.6, 106.2, 105.6, 65.9, 56.4, 46.1, 45.1, 36.6, 27.1, 16.6, 13.6. HRMS (ESMS) Calculated for C_38_H_45_N_13_O_6_ [M+H]^+^: 780.3694, found 780.3663.

**Synthesis of diABZI-V/C-DBCO (3):** To a stirred solution of (S)-2-((S)-2-amino-3-methylbutanamido)-N-(4-(hydroxymethyl)phenyl)-5-ureidopentanamide (400 mg, 1.05 mmol, 1eq) **(9)** and Hunig’s base (550 μL, 3.16 mmol, 3eq) in DMF 5 mL dropwise add solution of activated NHS ester (500 mg, 1.16 mmol, 1.1 eq) in DMF (2mL) stir overnight under argon atmosphere. The diethyl ether added to precipitate out the solid product to obtain the desired product with terminal benzyl alcohol as an intermediate **(11)** (702 mg, 1.10 mmol, 96%). To the stirred solution of intermediate alcohol (550 mg, 0.79 mmol, 1 eq) and Hunig’s base (689 μL, 3.95 mmol, 5 eq) in DMF (5 mL) add 4-nitrophenyl carbonochloridate (319 mg, 1.58 mmol, 2.0 eq) and stir overnight. The diethyl ether added to precipitate out the solid product to obtain as crude product. The crude solid was purified over silica gel chromatography (DCM:MeOH 0-25%) to obtain the de desired product (113 mg, 0.13 mmol,17%) **(1)**.

Compound **1** was reacted in a solution of diABZI-amine **(2)** (140 mg, 0.12 mmol, 1 eq) and Hunig’s base (0.11 mL, 0.62 mmol, 5 eq) in DMF (3 mL) stirred under argon atmosphere at room temperature. A solution of activated ester (113 mg, 0.13 mmol, 1.05 eq) in DMF (2 mL) was added dropwise to the reaction mixture. The reaction mixture was stirred for overnight. The solvent DMF was evaporated in vacuo to obtain the crude desired product. The crude solid was purified over silica gel chromatography (DCM:MeOH 0-25%) to obtain the desired product diABZI-V/C-DBCO **(3)** as an light pink solid (170 mg, 0.11 mmol, 86%). HRMS (ESMS) Calculated for C78H89N19O13 [M+Na]+: 1522.6785, found 1522.6821.

**Isothermal Calorimetry:** Recombinant hSTING was synthesized in Escherichia coli (New England Biolabs, T7 SHuffle Express line) and purified by affinity chromatography. Buffer exchange was performed prior to ITC (pH 7.5: PBST, 150 mM NaCl, 3 mM EDTA, 0.05% Tween 20), using Amicon Ultra 4 mL centrifugal filters (Millipore, Etobicoke, Canada). ITC experiments were performed on a TA Instruments Affinity ITC instrument. 24 total injections were performed using the following instrument settings: cell temperature 25°C, reference power 10 µCal/second, initial delay 240 seconds, stirring speed 75 rpm, feedback mode/gain high, and injection volume 2 µL for 10 seconds spaced at 120 second intervals with a filter period of 10 seconds. hSTING was set in the cell at a concentration of 10 µM and a volume of 350 µL. Agonists were prepared at a stock concentration of 20 mM in DMSO and diluted using pH 7.5 PBST to 150 µM for titration by the syringe (120 µL). Data were analyzed using TA Instruments NanoAnalyze Software.

**Cathepsin B Release Assay:** Recombinant Mouse Cathepsin B (RND Systems) was prepared at 100 µg/mL in MES buffer (pH 5.0) upon opening and kept in -80 °C conditions when not in use. To activate the enzyme, Cathepsin B was diluted to 10 µg/mL in MES buffer (pH 5.0) containing 5 mM DTT and incubated at 37 °C for 15 minutes. After activation, 0.5 µg/mL Cathepsin B was combined with 100 µM of substrate at 37 °C for 48 hours at a total volume of 100 µL. Activity was determined by observing molecular weight shifts in the substrate using matrix assisted laser desorption and ionization mass spectrometry (MALDI-MS).

**MALDI-MS:** 3 µL of Matrix (15 mg mL-1 THAP in dry acetone) was combined with 1 µL of sample from the Cathepsin B activity assay and spotted on a stainless steel MALDI-MS plate (Bruker). After evaporation of matrix, two technical replicates were collected for each spot using FlexControl software (Bruker Daltonics) on a Bruker AutoFlex MALDI-TOF. The laser pulse rate was 1000 Hz and spectra were obtained with a mass window of 400-4000 m/z at high resolution (4.00 GS/s). FlexAnalysis software (Bruker Daltonics) was used to obtain baseline spectra for all samples.

**THP1-Dual IFN-β Reporter Cell Assay:** THP1-Dual cells (InvivoGen) were cultured in Roswell Park Memorial Institute (RPMI) 1640 Medium (Gibco) supplemented with 2 mM L-glutamine, 25 mM HEPES, 10% heat-inactivated fetal bovine serum (HI-FBS; Gibco), 100 U ml^−1^ penicillin/100 μg ml^−1^ streptomycin (Gibco), and 100 µg/mL Normocin. Cells were subjected to 10 µg/mL Blasticidin and 100 µg/mL Zeocin for continual selection after every cell passage. 96-well plates (REF 655180; Greiner Bio-One) were used for screening agonist activity. Reporter cells were seeded at 25,000 cells/well in 100 µL media and treatments were administered in 100 µL of medium. Results were collected 24 hours after treatment using a Quanti-Luc™ (InvivoGen) assay on cell supernatants following manufacturer’s instructions. Luminescence was quantified using a plate reader (Synergy H1 Multi-Mode Microplate Reader; Biotek) after supernatants were transferred to opaque-bottom 96-well plates (REF 655073; Greiner Bio-One). EC_50_’s were determined using a variable slope non-linear regression fit in GraphPad Prism.

**Splenocyte Isolation and IFN-β ELISA:** Spleens were harvested from Female C57BL/6 mice (8 weeks old), mechanically disrupted into single-cell suspensions through a 70 μm cell strainer (FisherbrandTM; Thermo Fisher Scientific), and suspended in complete RPMI 1640 medium (Gibco) supplemented with 10% FBS, 10% HI-FBS (Gibco), 100 U ml−1 penicillin/100 μg ml−1 streptomycin (Gibco), 50 μM 2-mercaptoethanol, and 2 mM L-glutamine. The cells were centrifuged for 5 min at 1500 rpm and resuspended in ACK lysis buffer (KD Medical) for 5 minutes. Cells were centrifuged and resuspended in fresh media at a concentration of 3 million cells per mL. Cells were seeded in a 96 well round bottom plate (brand) at 100 μL per well and treatments were administered in 100 μL of medium. Results were collected 24 hours after treatment using a mouse IFN-β solid-phase sandwich ELISA kit (Invivogen Cat#42400-1) on cell supernatants following manufacturer’s instructions. Luminescence was quantified using a plate reader (Synergy H1 Multi-Mode Microplate Reader; Biotek).

**BMDM Isolation and IFN-β ELISA:** Bone marrow cells were harvested from femurs and tibias of 6–8 week-old female C57Bl/6 J mice by flushing them with culture media (RPMI 1640 medium supplemented with 10% heat-inactivated FBS, 100 U/mL penicillin, 100 μg/mL streptomycin, 2 mM L-glutamine, 10 mM HEPES, 1 mM sodium pyruvate, 1× nonessential amino acids, 50 μM β-mercaptoethanol, and 20 ng/mL M-CSF). The cell suspension was passed through a 70 μM cell strainer (FisherbrandTM; Thermo Fisher Scientific) and cells were centrifuged for 5 min at 1500 rpm and resuspended in 5 mL of ACK lysis buffer. After 5 minutes, 25 mL of PBS was added to the cell suspension and the cells were centrifuged at 1500 rpm for 5 min and resuspended in complete media (FisherbrandTM; Thermo Fisher Scientific) and seeded at 500,000 per mL in 100 × 15 mm non-tissue-culture-treated Petri dishes (Corning Inc.) to induce differentiation into BMDMs. Cell were cultivated in a humidified chamber maintained at 37 ◦C with 5% CO2 and media was changed every three days. After 8-10 days, BMDM phenotype was confirmed using flow cytometry (CD11b+F4/80+). BMDMs were were seeded in a 96 well round bottom plate at 200,000 cells per well in 100 μL per well and treatments were administered in 100 μL of medium. Results were collected 24 hours after treatment using a mouse IFN-β solid-phase sandwich ELISA kit (Invivogen Cat#42400-1) on cell supernatants following manufacturer’s instructions. Luminescence was quantified using a plate reader (Synergy H1 Multi-Mode Microplate Reader; Biotek).

**Polymer Synthesis and Characterization:** Reversible-addition-fragmentation (RAFT) polymerization was used to synthesize a series of poly(Dimethylacrylamide-co-azide-ethylmethacrylate) (poly(DMA-co-AzEMA)) polymers. A feed ratio of 93% DMA and 7% AzEMA was utilized with the chain transfer agent (ethylsulfanylthiocarbonyl)-sulfanylvpentanoic acid (ECT) and the initiator V70. A CTA-to-initiator ratio of 5 was utilized and the reaction was run in 100% dioxane with the mass fraction of CTA and monomer of 0.4. The reaction mixture was sealed and purged under argon gas for 20 minutes, and the reaction was run in a 40 degree C oil bath for 18 hours. Polymers were purified through acetone-to-water dialysis and dried via lyophilization. Through conversion NMR, molecular weights of 26338 and 111884 Da were calculated through the percent loss of vinyl peaks post polymerization (5.5-6.5 ppm) using DMA methyl peaks as reference (3 ppm). To determine composition (DMA:AzEMA), we compared the ratio of DMA peaks (3 ppm corresponding to 6 DMA protons) and AzEMA (4.1 ppm corresponding to 2 AzEMA protons post purification. A refractometer was utilized to obtain dn/dc values and applied in gel permeation chromatography (GPC) (EcoSEC Elite HLC-8420GPC). Copper-free click chemistry was utilized to conjugate Cyanine5-DBCO (Lumiprobe) to polymers using a 2:1 dye-to-polymer molar ratio in 100% DMSO. Reactions were continuously stirred at room temperature for 24 hours. Both platforms were purified via dialysis using 12-14kDa regenerated cellulose tubing (Spectra/Por) using a DMSO-to-molecular-grade water gradient and dried via lyophilization. Degree of labeling was determined through the absorbance at 650 nm (GENESYS 150, ThermoScientific).

**Drug Conjugation and Characterization via UV-Vis Spectrophotometry:** Copper-free click chemistry was utilized to conjugate diABZI-V/C-DBCO to the polymer platforms. Cy5-labeled polymers were dissolved at10 mg/ml in DMSO. A molar excess of 7:1 diABZI-V/C-DBCO to polymer was utilized for the 25kDa platform and a molar excess of 21:1 diABZI-V/C-DBCO to polymer was utilized for the 100kDa platform in 100% DMSO. Reactions were continuously stirred at room temperature for 24 hours. Both platforms were purified via dialysis using 12-14kDa regenerated cellulose tubing (Spectra/Por) using a DMSO-to-molecular-grade water gradient. 3kDa Amicon Ultra Centrifuge filters were used to concentrate conjugates. Each platform was further purified through Whatman 0.2 μM PTFE membrane syringe filters. Polymer concentration was determined using absorbance at 650 nm and drug concentration was determined at 325 nm with (GENESYS 150, ThermoScientific).

**BMDM Uptake:** Differentiated BMDMs were plated at 300,000 cells per well in a 96 well round bottom plate in 100 μL of previously described growth media. 25kDa conjugates or 100kDa conjugates were dosed at a total Cy5 concentration of 12.4 uM per sample and incubated at either 4 °C or 37 °C for 2 hours. After incubation, the plates were centrifuged at 1500 rpm for 5 minutes at 4 °C, and washed three times with refrigerated flow buffer (1% BSA in PBS). Cells were suspended in a 1% BSA solution containing 1:20000 of DAPI for a final cell suspension, at 100 µL per well. Data were collected and analyzed for cell uptake on a CellStream Flow Cytometer (Luminex) equipped with SSC, FFC, 405 (DAPI), and 642 (Cy5) nm lasers.

**BMDM Activation:** Differentiated BMDMs were plated at 300,000 cells per mL in 6 well tissue culture-treated plates in 2 mLs of previously described growth media. 25kDa conjugates or 100kDa conjugates were dosed at a total diABZI concentration of 1 uM per sample and incubated at 37 °C for 24 hours. After incubation, cells were collected using Cellstripper and plated at 500,000 cells per sample in 96 well round bottom plate in flow buffer (1% BSA in PBS). Samples were incubated with anti-mouse-CD86 and anti-mouse-MCHII (PE-Cy5 and BV605) for 1 hour. Cells were washed three times with refrigerated flow buffer and resuspended in a 1: 20,000 dilution of DAPI for a final cell suspension, at 100 µL per well. Data were collected and analyzed on a CellStream Flow Cytometer (Luminex) equipped with SSC, FFC, 405, 561, and 642 nm lasers.

**Blood Fluorescence Pharmacokinetics:** Healthy, 7 week old female C57BL/6 mice (The Jackson Laboratory) were injected with either PBS, 25kDa conjugate or 100kDa conjugate in PBS at a diABZI concentration of 0.012 μg/mouse in a 100 μL injection volume. Blood was sampled at various time points for 24 hour hours using heparinized capillary tubes (DWK Life Sciences). A 1:50 dilution of blood to PBS was centrifuged, and the plasma was collected for analysis. Cy5 concentration was determined by fluorescence intensity using a plate reader plate reader (Synergy H1 Multi-Mode Microplate Reader; Biotek), with an excitation wavelength of 645 nm and an emission wavelength of 675 nm after subtraction of PBS control. Pharmacokinetic analysis was performed in GraphPad Prism using a two-phase decay to determine elimination half-life.

**Tumor Models:** 6-8 week C57BL/6 femalemice (The Jackson Laboratory) were inoculated with the E0771 breast cancer model. 6-8 week Balb/C female mice (The Jackson Laboratory) were used for 4T1 breast cancer models. E0771 tumors were generated in models were inoculated at 3 x 10^5^ cancer cells per mouse in 100 µL injection volumes into left-side 4th mammary fat pad. Treatment was started when tumors reached ∼50-100 mm^3^ with the maximal endpoint at approximately 1500 mm^3^. 4T1 tumors were generated in models were inoculated at 1 x 10^6^ cancer cells per mouse in 100 µL injection volumes into left-side 4th mammary fat pad. Treatment was started when tumors reached ∼30 mm^3^ with the maximal endpoint at approximately 1500 mm^3^. Monotherapy treatments were administered through retroorbital injections of various drug concentration using 100 µL injection volumes in PBS. For combination therapy, I.P. injection of commercial anti-PD1 (BioXcell RMP1-14) were performed at 100 µg/mouse in 100 µL PBS injections.

**Organ and Tumor Biodistribution:** 7-week-old female C57BL/6 mice (The Jackson Laboratory) inoculated with 50-100mm^3^ E0771 mammary fat pad tumors were injected with either PBS, 25kDa conjugate or 100kDa conjugate in PBS at a diABZI concentration of 0.012 μg/mouse in a 100 μL injection volume. Mice were sacrificed 24 hours post injection. Lungs, livers, hearts, kidney, spleens, tumors, and contralateral mammary fat pad tissues were excised and washed in 1x PBS. Tissues were imaged the IVIS Lumina III (PerkinElmer). Fluorescence (radiant efficiency) was measured with a maximum value of 5.97 x 10^9^, and a minimum of 3.06 x 10^8^, and average radiant efficiency values (per cm2) were calculated using the Living Image software (version 4.5).

**TME Uptake:** Treatment was injected retro-orbitally, and at 24 hours following injection, mice were euthanized for analysis via flow cytometry. Tumors were harvested and dissociated using a Gentlemacs dissociator (Miltenyi Biotec). Following dissociation, tumors were treated with mouse tumor dissociation kit (Miltenyi Biotec) for 45 minutes at 37 °C shaking at 100 RPM. Samples were then dissociated again on a Gentlemacs dissociator and then mechanically dissociated through a 70 µm strainer to acquire a single-cell suspension. FC block to block non-specific binding took place for 15 minutes at 4 °C in the dark followed by surface stain for 30 minutes at 4 °C. Cells were washed and centrifuged at 380 rcf for 5 minutes then fixed in 2% paraformaldehyde for 10 minutes at room temperature. Samples were washed, resuspended, and run on a Cytek Aurora spectral flow cytometer (Cytek Biosciences) and then analyzed in FlowJo (BD Biosciences).

**Serum Cytokine ELISA:** Treatment was initiated at a tumor volume of approximately 50-100 mm^3^. Mice were treated with one dose of PBS, diABZI-V/C-DBCO, or 100kDa conjugates and blood was drawn 6 hours post systemic injection via cardiac puncture and collected in K2EDTA-coated tubes (BD Biosciences). Tubes were centrifuged at 1500 g for 15 min at 4 °C, and the serum was collected for analysis via IFN-β solid-phase sandwich ELISA kit (Invivogen Cat#42400-1).

**Tumor PCR:** Treatment was initiated at a tumor volume of approximately 50-100 mm^3^. Mice were treated with one dose of PBS, diABZI-V/C-DBCO, or 100kDa conjugates and tumors were harvested 6 hours post systemic injection. Tumors were lysed in RLT Plus lysis buffer using a TissueLyser II, (Qiagen). RNA was extracted with an RNeasy® Plus Mini Kit (Qiagen) according to the manufacturer’s protocol. An iScript cDNA synthesis kit (Bio-Rad) was used to synthesize cDNA per manufacturer’s protocol. RT-qPCR on the cDNA was run using TaqMan gene expression kits (primer and master mix) and run on the Bio-Rad CFX Connect Real-time System, with a threshold cycle number determination made by the Bio-Rad CFX manager software V.3.0. Primers: mouse Ifnb1 (Mm00439552_s1), mouse Tnf (Mm00443258_m1), mouse Cxcl10 (Mm00445235_m1), mouse Tmem173 (Mm01158117_m1), mouse IL-6 (Insert) and mouse Hmbs (Mm01143545_m1). Gene expression relative to Hmbs was calculated using 2^-(Cq-CqHmbs)^.

**Flow Cytometric TME Analysis:** Treatment was initiated at a tumor volume of approximately 50-100 mm^3^. For the myeloid panel, mice were treated with one dose of PBS, diABZI-V/C-DBCO, or 100kDa conjugates and tumors were harvested 24 hours post systemic injection for flow cytometry. For the T cell panel, mice were treated with three doses of PBS, diABZI-V/C-DBCO, or 100kDa conjugates, four days apart. Tumors were harvested 24 hours post final systemic injection for flow cytometry. Single-cell suspensions were prepared as described in uptake studies. FC block was applied for 15 minutes at 4 °C followed by surface stain for 30 minutes at 4 °C. Cells were fixed in Foxp3 Transcription Factor Staining Buffer (ebioscience) for one hour at room temperature then rinsed in permeabilization buffer (ebioscience). Intracellular staining took place for 30 minutes at room temperature in permeabilization buffer. Cells were rinsed, resuspended, and run on a Cytek Aurora spectral flow cytometer (Cytek Biosciences). Data analysis took place in FlowJo (BD Biosciences).

**Antibodies:** CD4 (RM4-5, BV605, Biolegend), CD44 (IM7, PerCP, Biolegend), CD366/Tim3 (RMT3-23, PE-Dazzle594, Biolegend), CD223/LAG3 (C9B7W, BV785, Biolegend), CD279/PD1 (29F.1A12, BV510, Biolegend), CD8α (53-6.7, AF488, Biolegend), CD69 (H1.2F3, PE-Cy7, Biolegend), CD62L (MEL-14, BV711, Biolegend) Ki-67 (SolA15, AF532, ebioscience), Granzyme B (NGZB, PE-Cy5.5, ebioscience), KLRG1 (2F1, sb645, ebioscience), TCR-β (S33-966, ef450, ebioscience), and FoxP3 (FJK-16S, PE, ebioscience). TCRb(H57-597, eFluor 450, eBioscience), CD4 (RM4-5, SB780, eBioscience), CD8a (53-6.7, BV605, BioLegend), CD11b (M1/70, BV510, BioLegend), CD11c (N418, BV711, BioLegend), GR-1 (RB6-8C5, PE/Cy7, eBioscience), F4/80 (BM8, AF488, BioLegend), and CD19 (6D5, PE, BioLegend).

**Blood Toxicity and Organ Pathology:** Healthy, 7 week old female C57BL/6 mice (The Jackson Laboratory) were injected (RO) with either PBS, or SAPCon[100kDa] in PBS at a diABZI concentration of 0.008 μg/mouse in a 100 μL injection volume for three total doses, four days apart. Mice were sacrificed 1 week post final treatment. Blood was harvested via cardiac puncture and used to prepare serum by centrifugation at 1500 g. The serum was tested by the Vanderbilt Translational Pathology Shared Resource for various protein levels that would be indicative of potential liver and kidney toxicity. Livers were harvested, fixed in a 10% formalin in PBS solution, paraffin embedded and sectioned for haematoxylin and eosin staining. Interpretation was completed by a board-certified veterinary pathologist at Vanderbilt University Medical Center.

**Statistics:** All data were plotted and analyzed using GraphPad Prism 10 and reported as mean +/-SD or SEM. Grubb’s test was utlilized to identify outliers. Comparisons between two groups utilized an unpaired Student’s T-test. A one-way ANOVA with post-hoc Tukey’s correction was used for multiple comparisons. Tumor volume significance was identified through a two-way ANOVA followed by a Tukey’s adjustment for multiple comparison. Surivial curve comparisons were make using a Log-rank test.

## Acknowledgements

The authors thank C. Duvall for use of IVIS Imaging System. We thank the core facilities of the VUMC Flow Cytometry Shared Resource, supported by the Vanderbilt Digestive Disease Research Center (DK058404) and the Vanderbilt Ingram Cancer Center (P30 CA68485), the Vanderbilt Institute of Nanoscale Sciences and Engineering (VINSE), the Vanderbilt University Small Molecular NMR Facility, and Translational Pathology Shared Resource supported by NCI/NIH Cancer Center Support Grant P30CA068485. This research was supported by grants from the Susan G. Komen (CCR19609205 to JTW), The National Institutes of Health (R01 CA245134 to JTW), the National Science Foundation (CBET-1554623 to JTW), a Vanderbilt Ingram Cancer Center (VICC) Ambassador Discovery Grant (JTW), VICC Support Grant (P30 CA068485), and funds provided by the Vanderbilt University School of Engineering (JTW). BRK acknowledges postdoctoral funding support from the PhRMA Foundation Postdoctoral Fellowship in Drug Delivery. AJK was supported by the NIH Microenvironmental Influences in Cancer Training Grant (T32CA009592). TLS acknowledge funding support through the National Science Foundation Graduate Research Fellowship Program under grant number 193793. Any opinions, findings, and conclusions or recommendations expressed in this material are those of the author(s) and do not necessarily reflect the views of the National Science Foundation. Schematics were made using Biorender.com.

