## Supporting Information for "STING-Activating Polymer-Drug Conjugates for Cancer Immunotherapy"

### Supplemental Data:

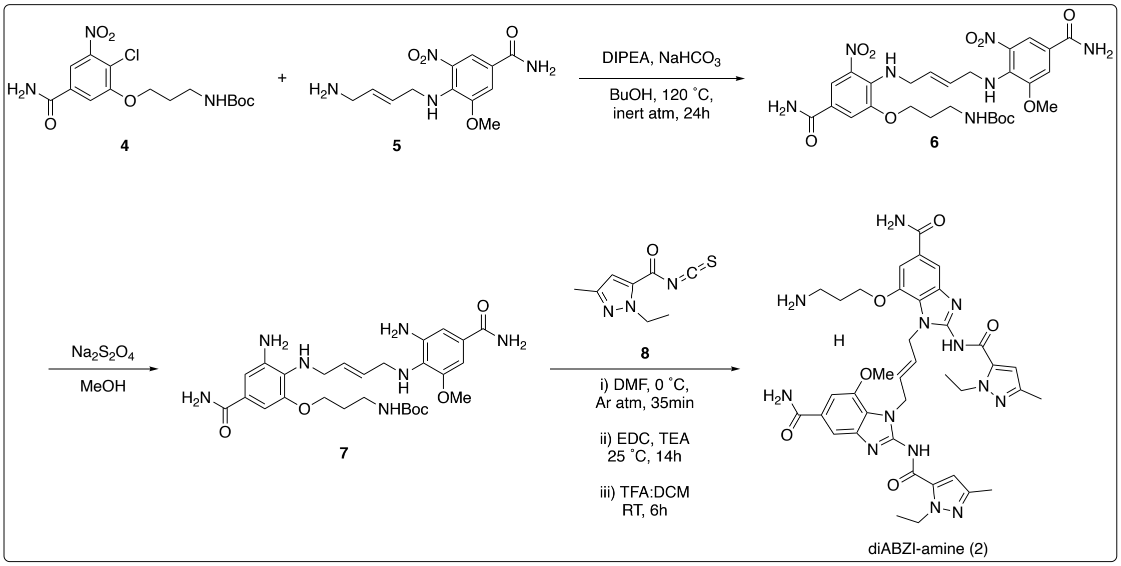


**Scheme S1:** Synthesis of diABZI-Amine

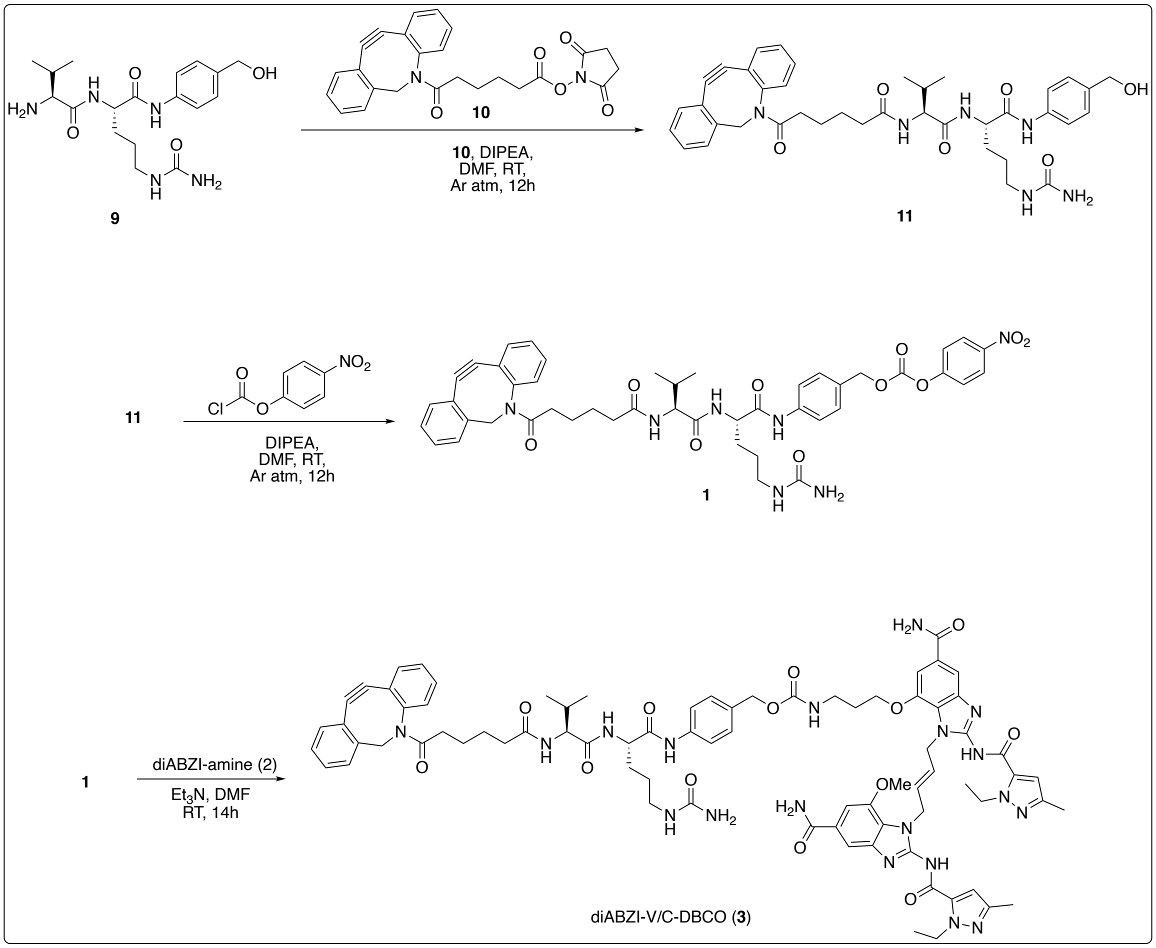


**Scheme S2:** Synthesis of diABZI-V/C-DBCO

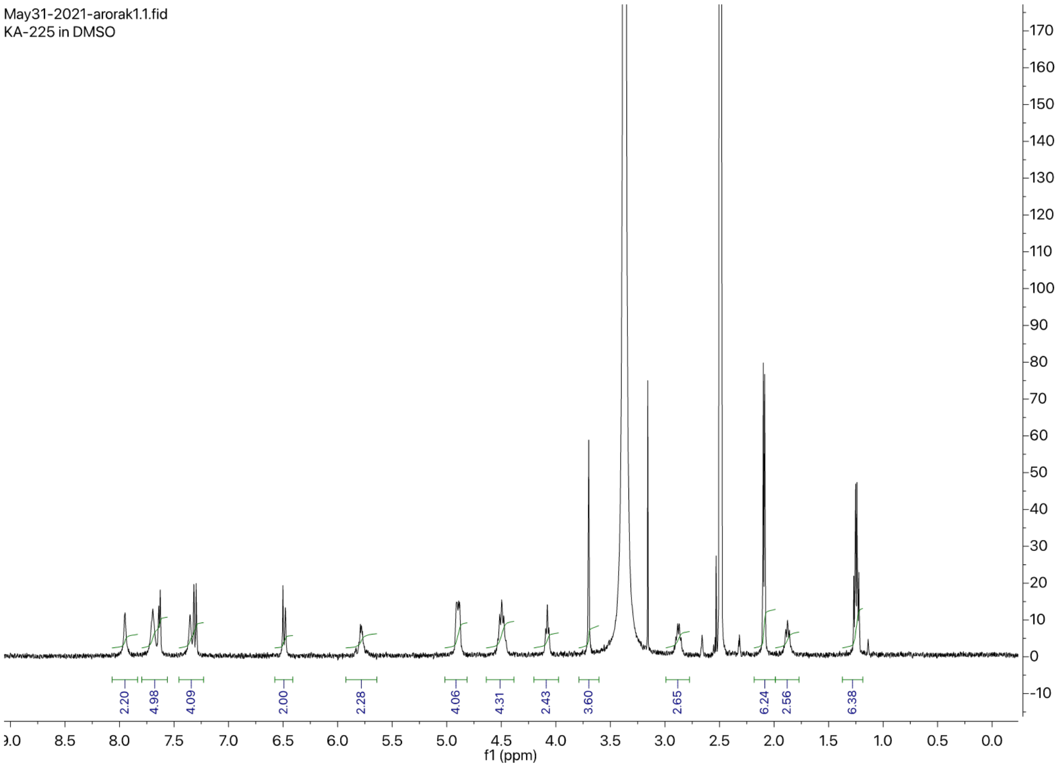


**Figure S3:** H[1] NMR of diABZI-Amine

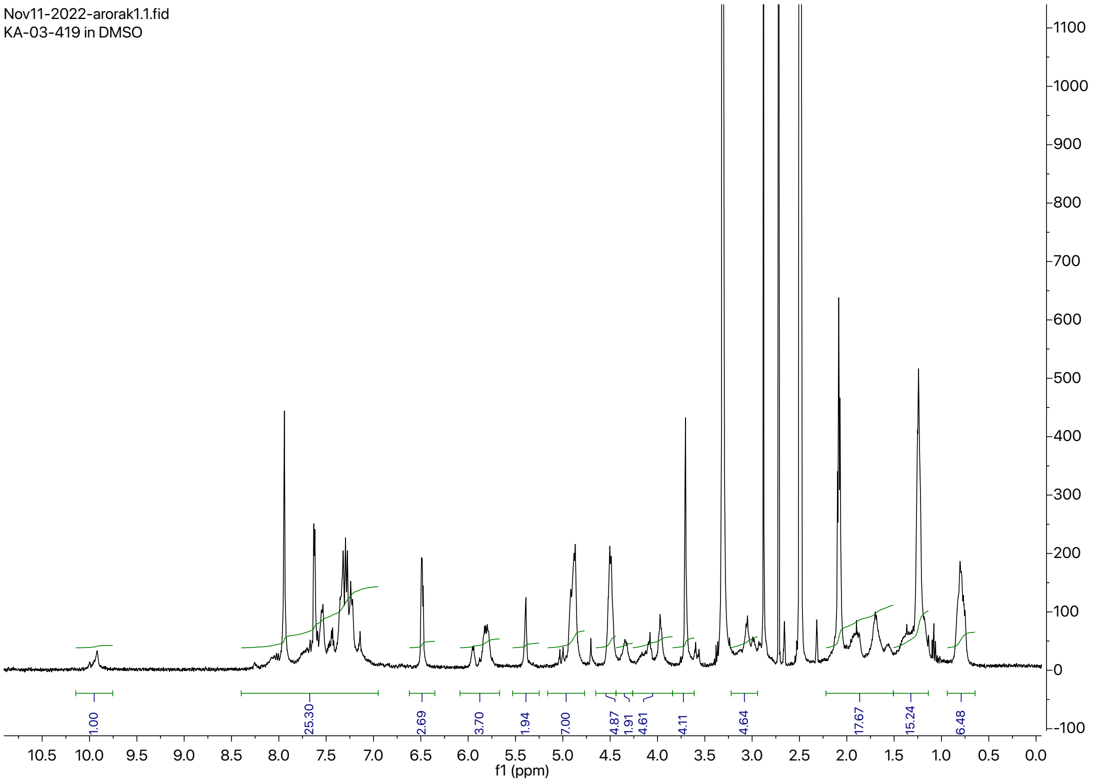


**Figure S4:** H[1] NMR of diABZI-V/C-DBCO

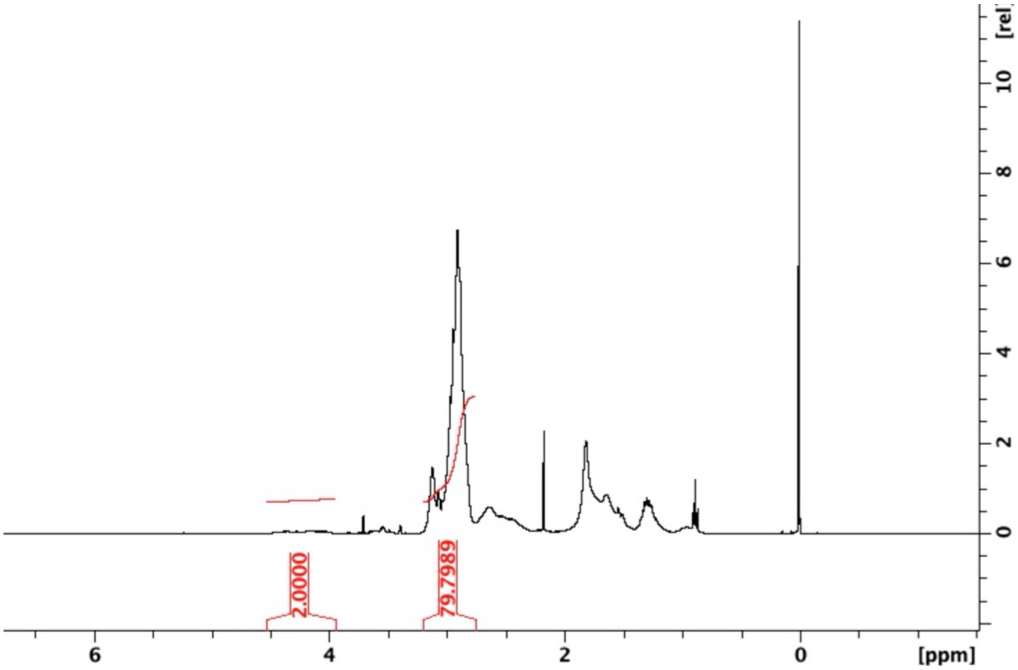

**Figure S5:** $H^1$ NMR of 25kDa DMA-co-AzEMA

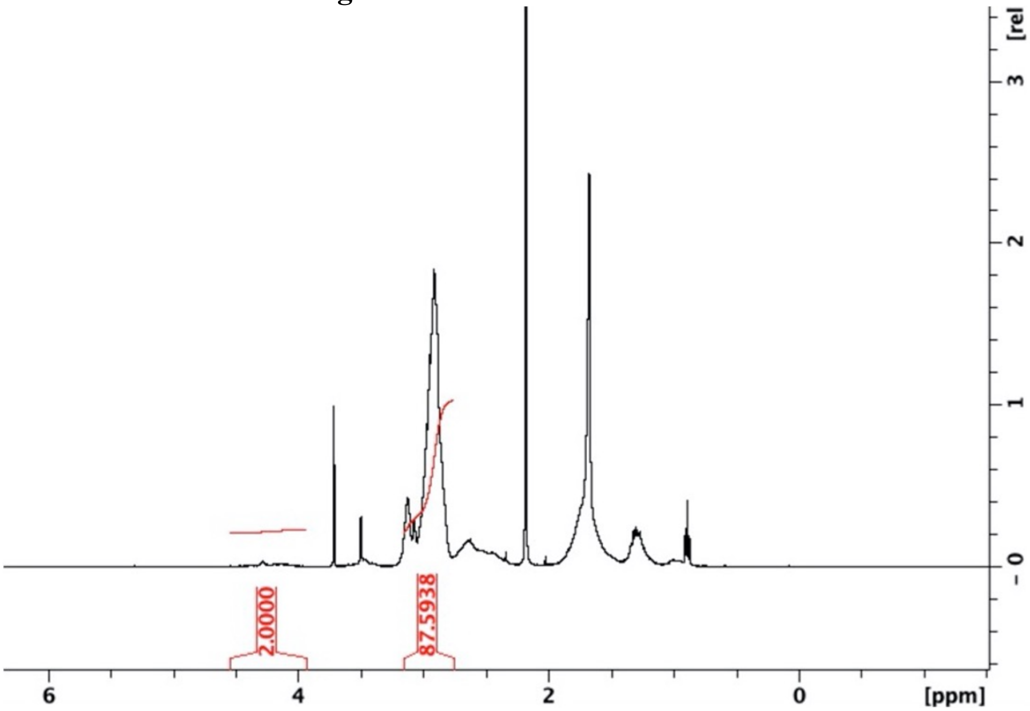


**Figure S6:** $H^1$ NMR of 100kDa DMA-co-AzEMA

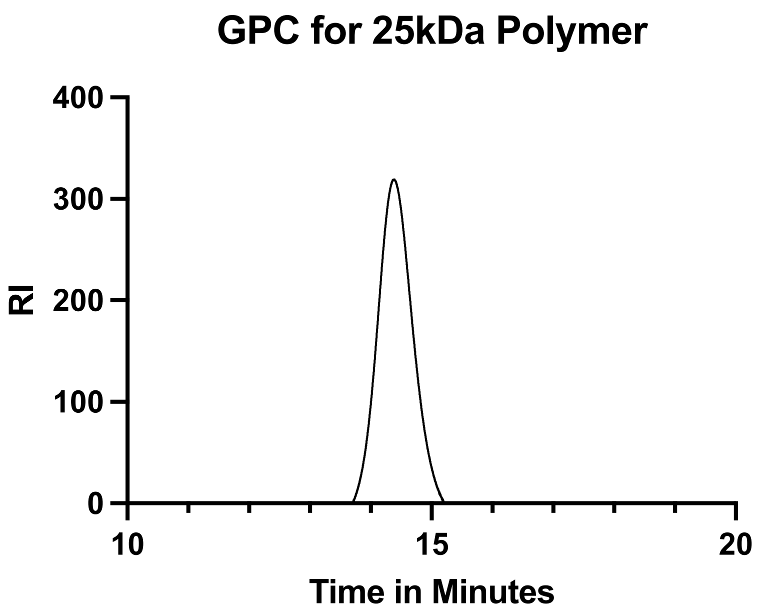


**Figure S7:** GPC of 25kDa DMA-coAzEMA

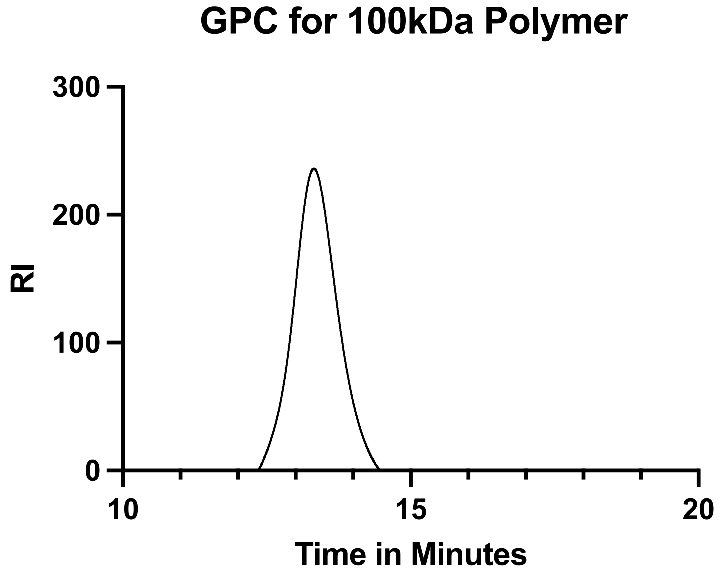


**Figure S8:** GPC of 100kDa DMA-coAzEMA

| Polymer | Mw NMR | Mw GPC | Mn GPC | PDI | Azides/chain (approximate) |
| --- | --- | --- | --- | --- | --- |
| **25kDa** | **25,247** | **28,357** | **27,483** | **1.03** | **19** |
| **100kDa** | **100,559** | **131,751** | **116,393** | **1.13** | **72** |

**Table S9:** Polymer Characterization

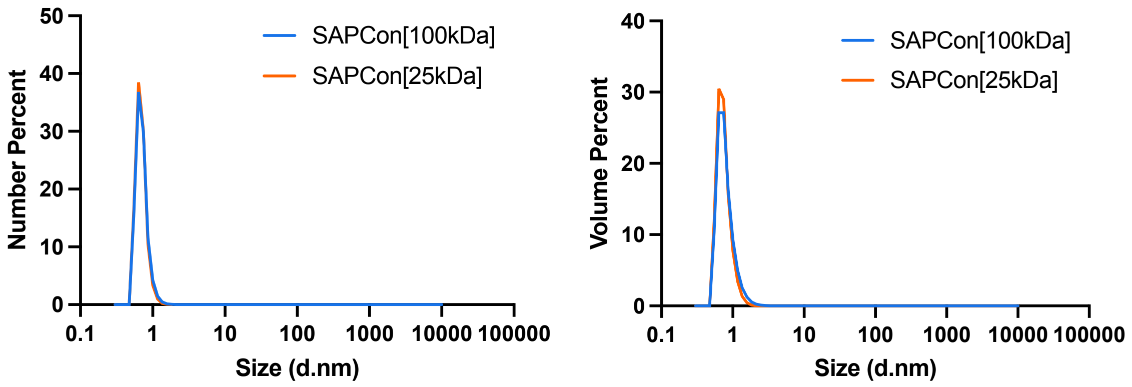


**Figure S10:** Dynamic light scattering of SAPCon[25kDa] and SAPCon[100kDa]

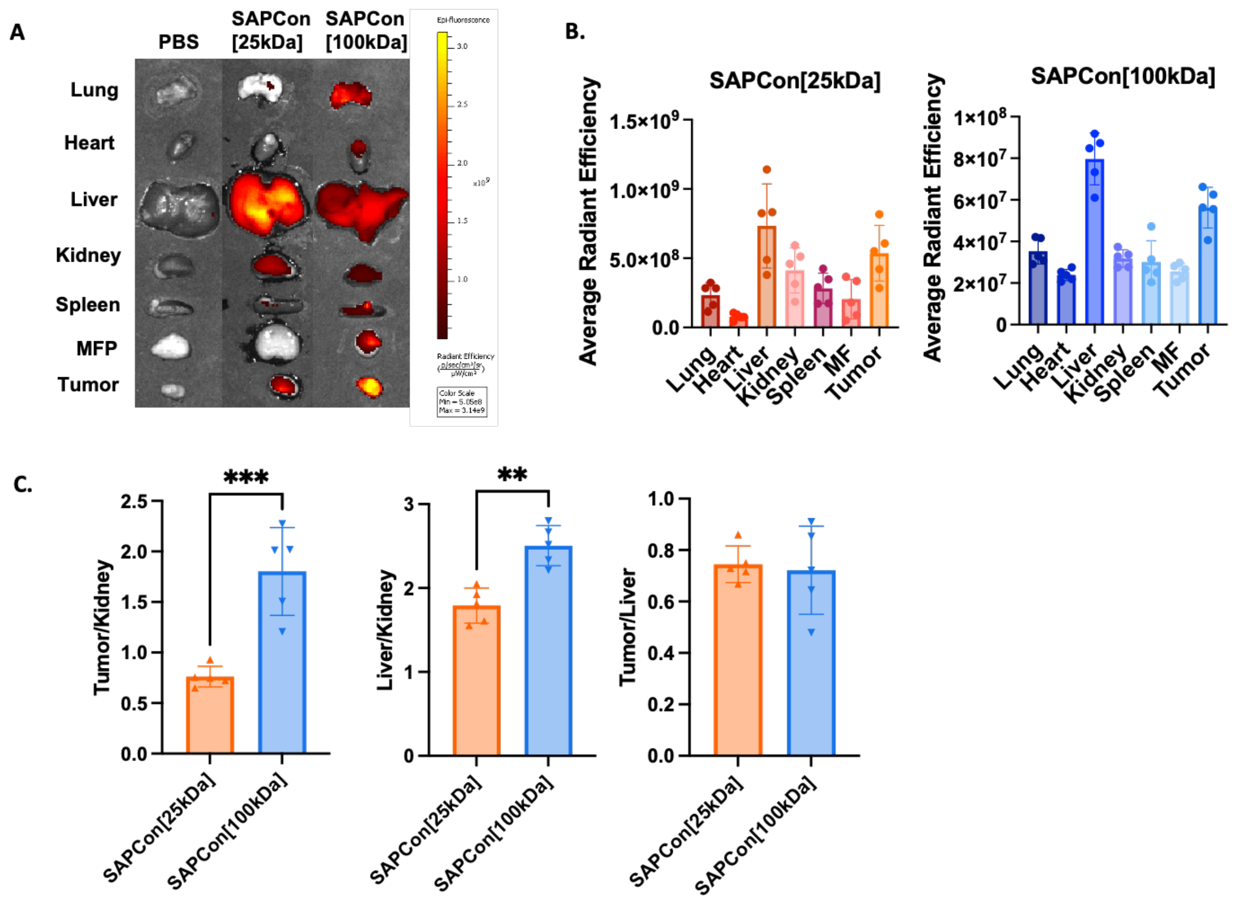


**Figure S11:** Biodistribution at 24 hours. **A)** Representative IVIS fluorescent images of excised EO771 tumors and organs 24 h following administration of Cy5-labeled SAPCon. **B)** Quantification of tissue fluorescence measured with IVIS imaging 24 h following administration of Cy5-labeled SAPCon (n=5). **C)** Ratios of tissue fluorescence between tumor and/or organs.

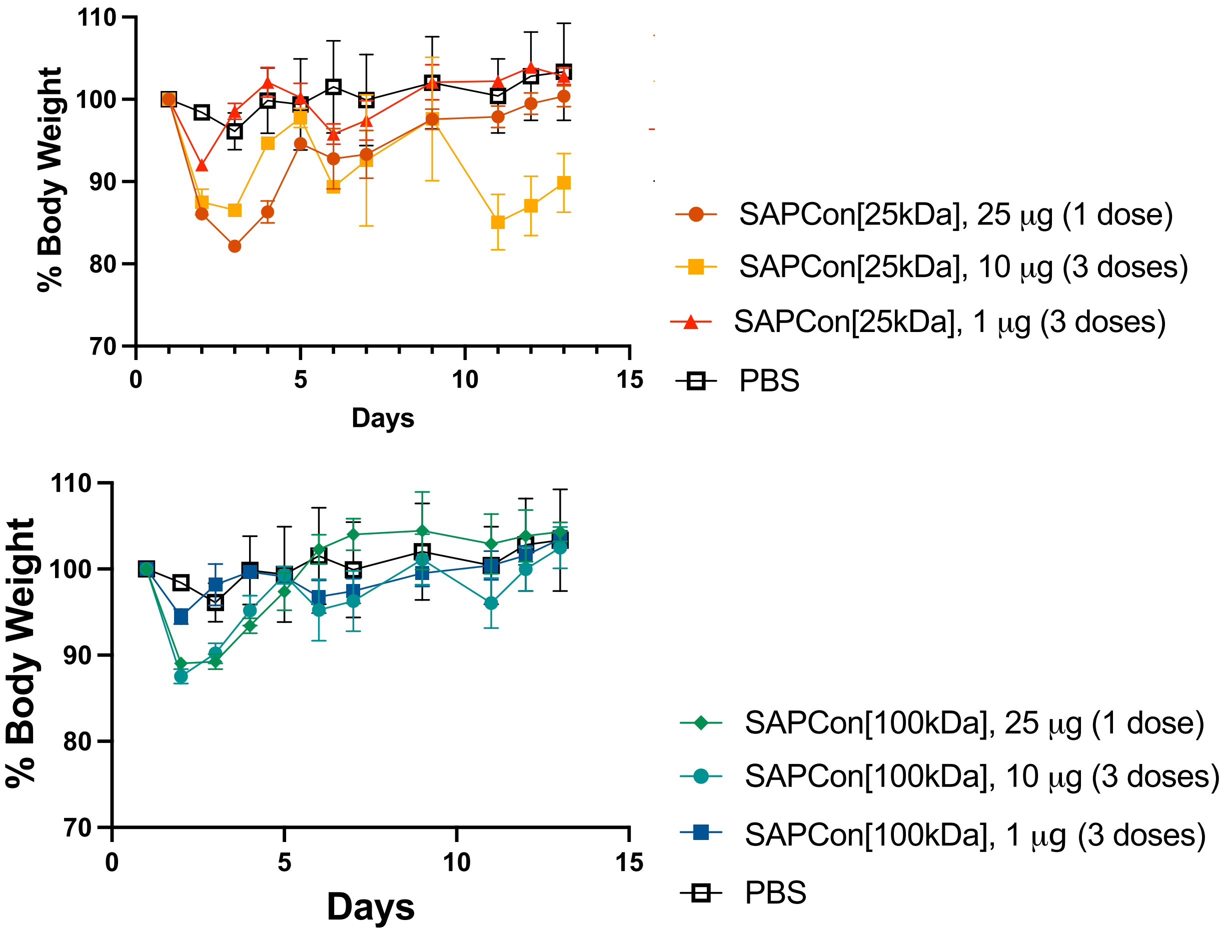


**Figure S12**: Transient weight loss in dose-finding toxicity study for SAPCon[25kDa] and SAPCon[100kDa] in healthy (n=13) C57BL/6 mice.

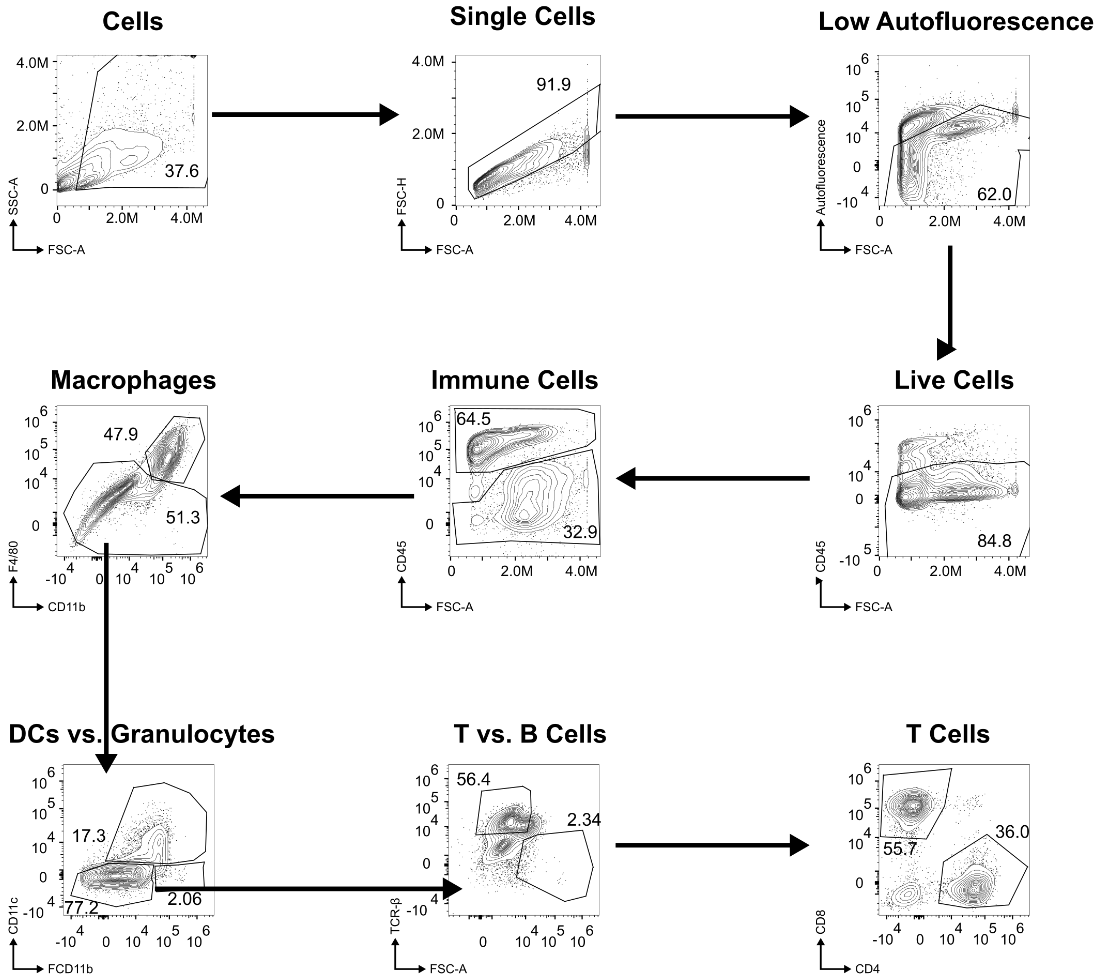


**Figure S13:** Flow cytometry gating for cell identification in Cy5 tumor uptake panel

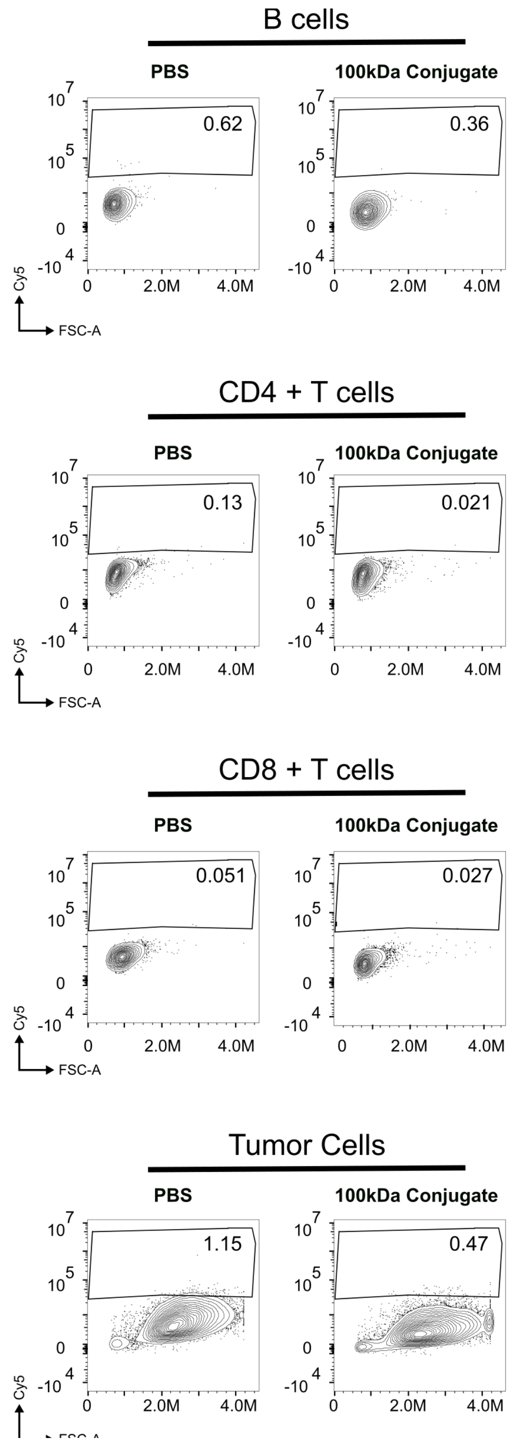


**S14:** Flow cytometry gating for Cy5 uptake in other cell populations within the tumor

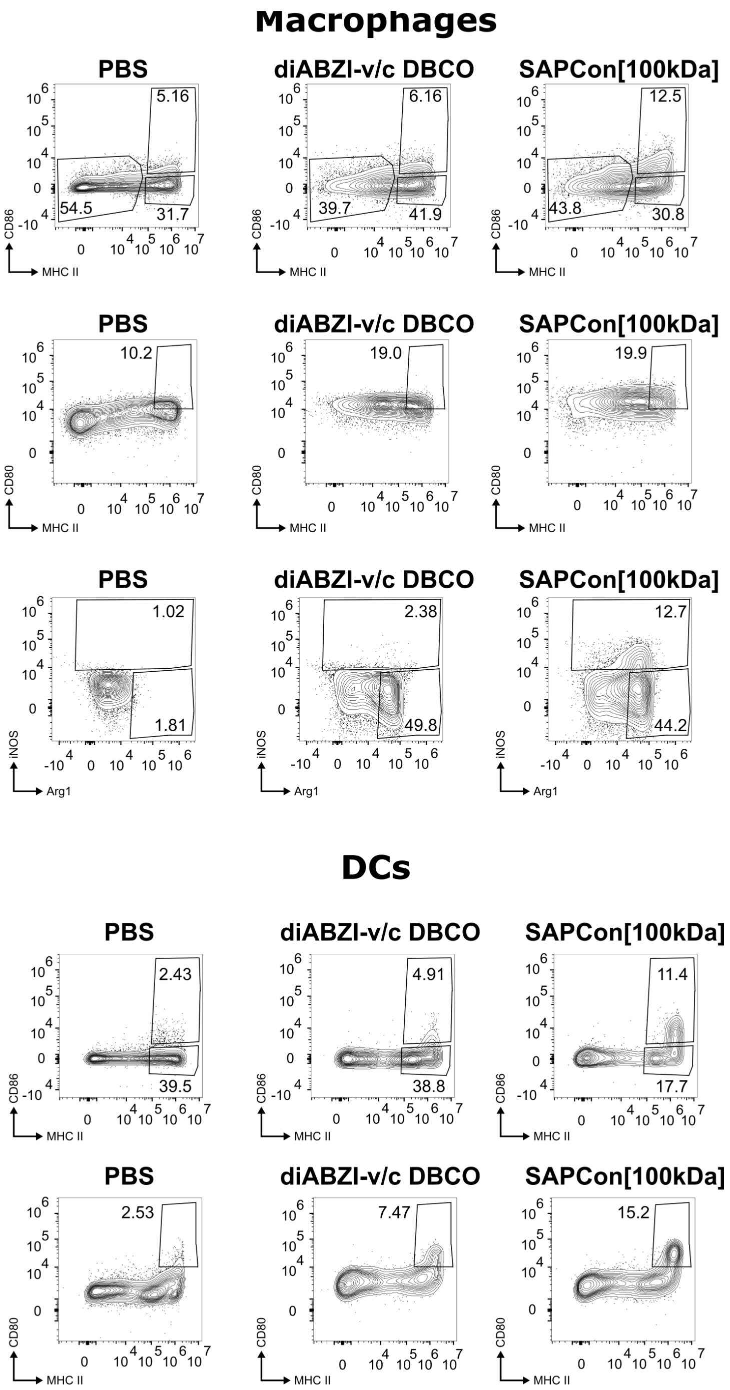


**Figure S15:** Gating scheme for macrophage and dendritic cell activation within the tumor

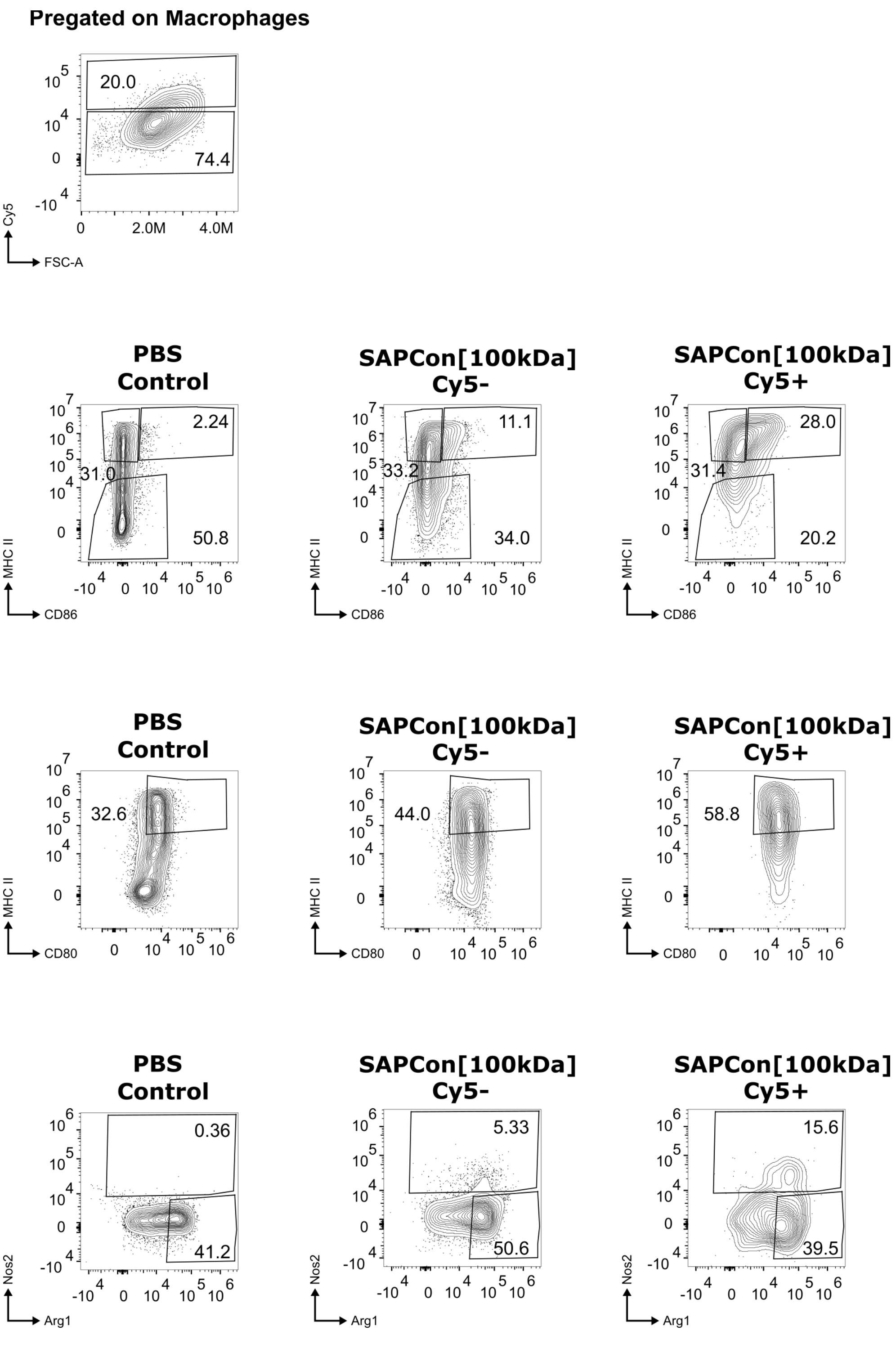


**Figure S16:** Gating scheme for Cy5 uptake and macrophage activation within the tumor

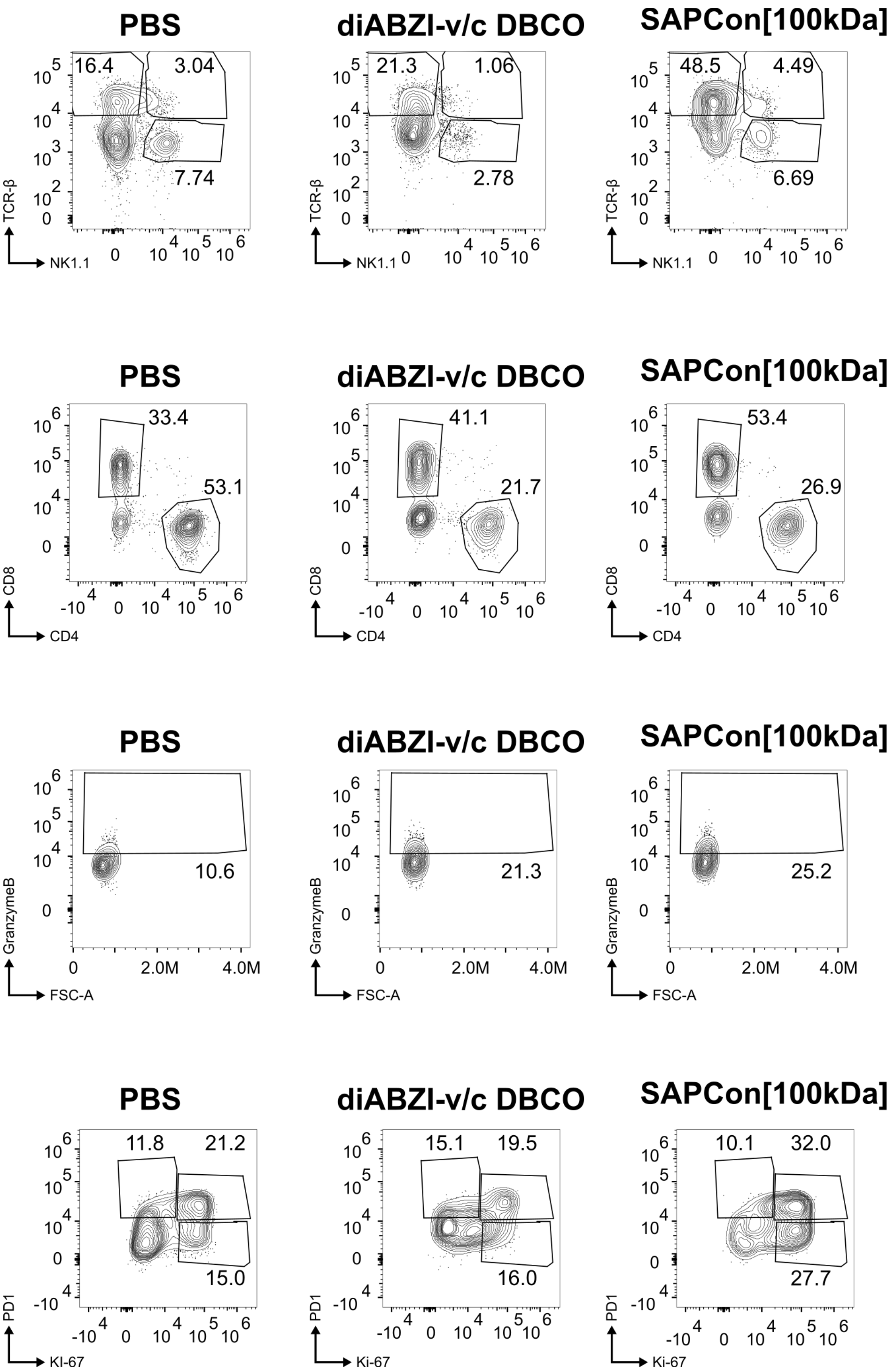


**Figure S17:** Gating scheme for T cell identification within the tumor

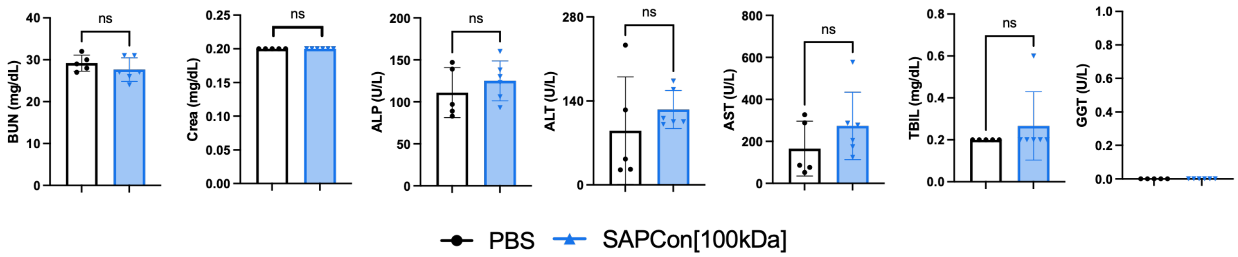


**Figure S18:** Blood toxicity marker analysis post treatment in healthy C57BL/6 mice (n=6)

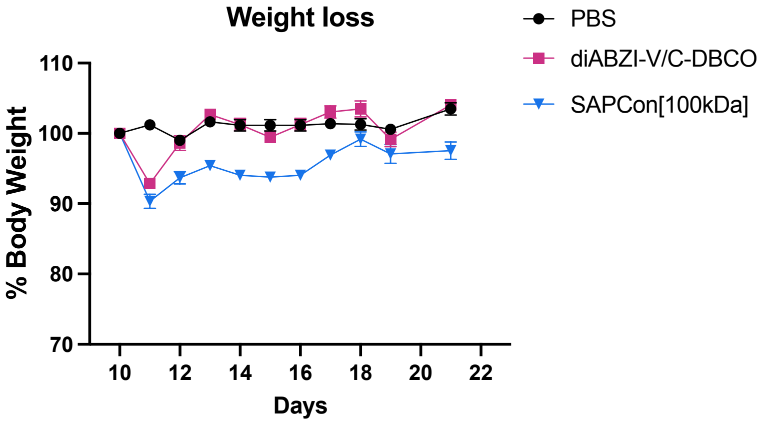


**Figure S19:** Mouse weight loss in EO771 therapy study

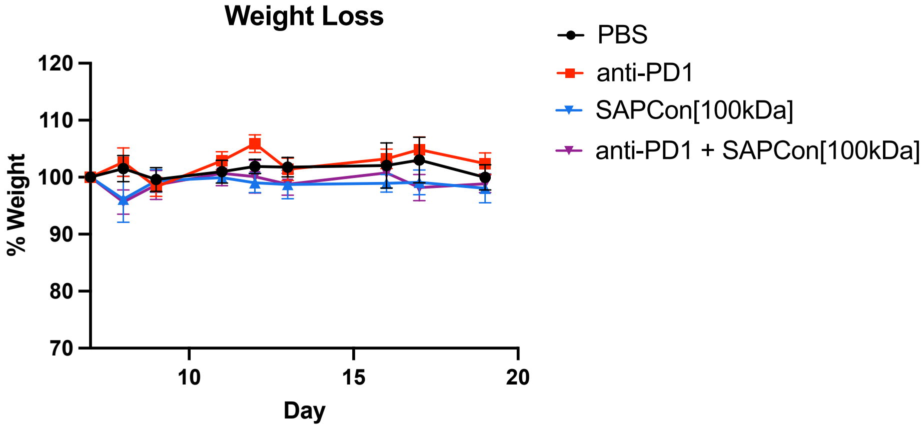


**Figure S20:** Mouse weight Lloss in 4T1 therapy study
